# Spectral Cluster Supertree: fast and statistically robust merging of rooted phylogenetic trees

**DOI:** 10.1101/2024.05.07.593083

**Authors:** Robert N. McArthur, Ahad N. Zehmakan, Michael A. Charleston, Gavin Huttley

## Abstract

The algorithms for phylogenetic reconstruction are central to computational molecular evolution. The relentless pace of data acquisition has exposed their poor scalability and the conclusion that the conventional application of these methods is impractical and not justifiable from an energy usage perspective. Furthermore, the drive to improve the statistical performance of phylogenetic methods produces increasingly parameter-rich models of sequence evolution, which worsens the computational performance. Established theoretical and algorithmic results identify supertree methods as critical to divide-and-conquer strategies for improving scalability of phylogenetic reconstruction. Of particular importance is the ability to explicitly accommodating rooted topologies. These can arise from the more biologically plausible non-stationary models of sequence evolution.

We make a contribution to addressing this challenge with Spectral Cluster Supertree, a novel supertree method for merging a set of overlapping rooted phylogenetic trees. It offers significant improvements over Min-Cut supertree and previous state-of-the-art methods in terms of both time complexity and overall topological accuracy, particularly for problems of large size. We perform comparisons against Min-Cut supertree and Bad Clade Deletion. Leveraging two tree topology distance metrics, we demonstrate that while Bad Clade Deletion generates more correct clades in its resulting supertree, Spectral Cluster Supertree’s generated tree is generally more topologically close to the true model tree. Over large datasets containing 10000 taxa and ~500 source trees, where Bad Clade Deletion usually takes ~2 hours to run, our method generates a supertree in on average 20 seconds. Spectral Cluster Supertree is released under an open source license and is available on the python package index as sc-supertree.

This research was undertaken with the assistance of resources and services from the National Computational Infrastructure (NCI), which is supported by the Australian Government.

## 1 Introduction

The relentless pace of DNA sequence data acquisition has exposed the poor scalability of phylogenetic reconstruction algorithms, establishing that new scalable and accurate algorithms are required for phylogenetic analysis. The well known sensitivity of phylogenetic results to data features that violate models of sequence evolution is increasing the interest in applying non-stationary models to phylogenetic inference (e.g. [5, 11, 29]) including estimation of rooted phylogenies [10, 34]. However, such models increase the number of free parameters and thus worsen computational performance. Theoretical results confirming the statistical consistency of the divide-and-conquer Disk-Covering Method (DCM) [9] identified this as a promising candidate for overcoming the robustness versus performance tradeoff, illustrated more recently by DACTAL [16]. DCM algorithms fundamentally rely on merging phylogenies estimated from overlapping subsets of the full data. While most prior DCM work has been focussed on unrooted trees, rooted topologies can also be handled. Given these considerations, we have sought to resolve one important bottleneck affecting the generalisation of DCM by developing a rooted supertree method that is statistically robust and scalable.

In this work, we introduce Spectral Cluster Supertree (SCS) for the case of rooted trees. SCS is a method which recursively partitions the set of all taxa in the source trees until a rooted supertree is generated. SCS derives its origins from Min-Cut Supertree [26], replacing the min-cut step with a substantially more scalable spectral clustering approach [27] to partition an internal graph. We also extend the method to make use of information such as branch lengths or depths of taxa in the source trees. These modifications lead to a much more efficient and accurate method, capable of solving problems with hundreds of source trees and ten thousand taxa in the order of seconds.

### 1.1 Related Work

Fleischauer and Böcker [7] performed an extensive systematic comparison among supertree methods in their paper that introduced Bad Clade Deletion (BCD). The comparison evaluated both rooted and unrooted supertree methods in terms of running time and accuracy of the generated supertree. Accuracy over generated datasets was measured by counting the number of splits of the generated supertree that were “true positives” (splits in the generated supertree that should occur), “false positives” (splits in the generated supertree that should not occur) and “false negatives” (splits in the generated supertree that should occur but do not). These values can be used to calculate the well-known F_1_ score.

The methods that were evaluated by the comparison included the Greedy Strict Consensus Merger (GSCM) [25]; FastRFS [32]; Matrix Representation with Parsimony (MRP) [1, 21]; SuperFine [31]; Combined Analysis using Maximum Likelihood (CA-ML) using RAxML [28]; and their own method BCD [7]. A rooted variant of GSCM exists that generates a supertree only containing clades that are compatible with all of the source trees [6]. FastRFS [32] generates an unrooted supertree optimising the Robinson-Foulds distance [23] to the source trees under a constrained search space using dynamic programming. MRP generates a rooted supertree by performing parsimony analysis on a Baum-Ragan encoding of the source trees [1, 21]. Superfine [31] is a meta-method which combines the Strict Consensus Merger [9] to merge the source trees together, with another supertree method (MRP yielded the best performance) to resolve polytomies; it generates unrooted trees. CA-ML concatenates the sequences used to generate each of the source trees and creates a supertree using maximum likelihood analysis; it has the potential to generate rooted trees depending on the chosen substitution model (for instance, a strand-symmetric model with IQ-TREE [15]).

Of the methods evaluated, CA-ML consistently performed the best in terms of topological accuracy of the constructed trees according to the F_1_ score, generally followed by BCD. However, BCD is much faster than CA-ML - on a particular dataset usually taking under eight seconds whereas CA-ML took most frequently approximately three days. On this dataset, all other methods usually took less than a minute to resolve a supertree with the exception of MRP which usually took between 15 minutes and an hour. BCD was consistently the fastest. The exact ordering of the accuracy of BCD and the other methods depended on the exact dataset, although BCD was most consistently at the top. Of particular importance when dealing with scalability was BCD’s performance on a large dataset containing an average of 5500 taxa. BCD with an option enabling branch length weighting performed the best in terms of accuracy here, as well as time taking 4-8 hours excluding GSCM (which exhibited very poor accuracy). The next fastest non-BCD algorithm was FastRFS usually taking 8-16 hours but also had poor accuracy in comparison. Superfine had the second best non-BCD accuracy and typically took between 16 hours and 3 days to resolve. MRP did not terminate within 14 days and on average exhibited worse accuracy than Superfine.

Spectral Cluster Supertree was created out of the desire for a rooted supertree algorithm that was capable of resolving a supertree containing several thousand of taxa in both an accurate and fast manner. Of the rooted supertree algorithms, based on the comparisons by Fleischauer and Böcker [7], BCD is clearly the most accurate and efficient. *We thus restrict our main comparisons in this paper to be between Spectral Cluster Supertree, and Bad Clade Deletion*.

The BCD [7] algorithm seeks to minimise the number of deleted characters in a Baum-Ragan [1, 21] matrix encoding of the source trees so that a consistent supertree is formed. In the matrix, rows represent the taxa, and columns represent the clades, or equivalently internal nodes, of a phylogenetic tree. An entry in the matrix is “1” if the taxon is in the clade, “0” if it is in the tree for the clade but not in the clade, and “?” otherwise. BCD aims to delete a minimum number of columns from this matrix to yield a matrix that defines a supertree.

From the matrix representation, BCD constructs a graph with an edge connecting clade *i* to taxon *j* if the corresponding entry in the matrix is 1. Any clades with no 0 entries in their column contain no useful information for the algorithm and are ignored in this process. If the graph is disconnected, the algorithm recurses over the separate components of the graph. The taxa in each of these components belong to different sides of the root in the tree. If the graph is not disconnected, BCD attempts to delete a subset of clades from the graph to make it so. The algorithm partitions the taxa over a minimum-weight cut of a transformation of the graph, only allowing clades to be deleted. The algorithm recurses over the components of the partition until a complete supertree is formed.

BCD also introduces a number of additional strategies to improve the accuracy of the supertrees and the time it takes to construct them. This includes a number of weighting strategies for the clades during the min-cut step, which can take into account information including branch lengths and bootstrap values. There is also a preprocessing step, where a rooted variant of the Greedy Strict Consensus Merger algorithm is used to collect clades that do not contradict with any of the source trees and ensure they are never cut [6, 25]. The algorithm also reduces the problem size by merging identical clades in the matrix representation. Parallelisation is additionally exploited when finding min-cuts and over the different partitions of the problem.

### 1.2 Using more Statistically Robust Distance Measures

It is conventional to use the Robinson-Foulds distance measure [23] to compare tree topologies. In the case of two rooted trees, it is determined from the number of clades that do not match between them. The Robinson-Foulds measure is known to exhibit poor statistical behaviour [2, 13], for instance, moving a single leaf in a caterpillar tree can maximise the metric [13]. We will later show in the materials and methods a mapping between the Robinson-Foulds distance, and the F_1_ score used by Fleischauer and Böcker [7]. Alongside the Robinson-Foulds distance, we also included the Matching Cluster distance [3]. Unlike the Robinson-Foulds distance, the Matching Cluster distance also considers the dissimilarity of clades that do not match. By capturing more detail of how topologies differ, it exhibits more robust statistical behaviour.

## 2 Definitions and Methods

### 2.1 Preliminaries

A *phylogenetic tree, T* = (*V, E*), is a connected, acyclic graph displaying evolutionary relationships over a set of *taxa, S*(*T*) - the leaves of the graph. Here, *V* is the set of leaves and internal vertices, and *E* is the set of edges that connect them. A tree can be *rooted* at an internal vertex to provide an orientation to the tree. In a *rooted bifurcating tree*, every internal vertex of *T* has degree three except for a special vertex of degree two that is labelled the root. A rooted tree is *multifurcating* if the degree of the root is greater than or equal to two and every other internal vertex of *T* is greater than or equal to three.

A taxon, *t* ∈ *S*(*T*), is said to be a *descendant* of an internal vertex, *v* ∈ *V*, of a phylogenetic tree if the path that connects the root to *t* passes through *v*. A *clade, C*, is the set of all descendants of an internal vertex. A clade is called *trivial* if |*C*| = 1 (it is a single leaf) or *C* = *S*(*T*) (the clade for the root of the tree). All other clades are *non-trivial*. Two taxa *u, v* ∈ *S*(*T*) are said to belong to a *proper cluster* of *T* if the unique path connecting *u* to *v* does not pass through the root [26]. That is, two taxa belong to a proper cluster if they are both descendants of an identical non-root vertex of *T* .

For a graph *G* = (*V, E*), a *contraction* of two vertices *u, v* ∈ *V* where (*u, v*) ∈ *E* gives a new graph *G*^*′*^ = (*V* ^*′*^, *E*^*′*^) formed by removing the edge from *E* and combining *u* and *v* into a single vertex, deleting any parallel edges. For contraction in the context of a weighted graph, all parallel edges except the one of maximal weight are deleted.

We say that a tree *A* is a *subtree* of another tree *B* if *A* can be obtained from *B* by deleting all taxa not in *A* from *B* and performing a sequence of contractions. If *A* is a subtree of *B*, then we say that *B displays A*. A collection of trees is called *compatible* if there exists a phylogenetic tree that displays all of them.

Let *T* be a phylogenetic tree and *X* be a set of taxa. The *induced subtree T* |_*X*_ is the maximally sized subtree of

*T* such that *S*(*T* |_*X*_) ⊆ *X*. A multiset of trees 𝒯 = {*T*_1_, *T*_2_, …, *T*_*n*_} can be induced on *X* such that 𝒯 |_*X*_ = {*T*_1_|_*X*_, *T*_2_|_*X*_, …, *T*_*n*_|_*X*_ }. If inducing a subtree would remove all taxa in a tree, it does not appear in the resulting set.

A rooted supertree algorithm is one which takes as its input a multiset of rooted trees 𝒯, called *source trees*, and returns a single rooted tree *T* such that 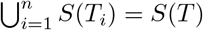.

### 2.2 Min-Cut Supertree

Min-Cut Supertree is a well-studied polynomial-time algorithm for merging rooted phylogenetic trees [26]. It works by recursively partitioning the taxa until a rooted tree is formed. There are proven properties of the algorithm, including that if all of the source trees are compatible, then the returned tree displays all of them, and any triple that is displayed by all of the source trees is displayed in the supertree.

#### 2.2.1 Proper Cluster Graph

Given a multiset of rooted source trees 𝒯 = {*T*_1_, *T*_2_, …, *T*_*n*_} and an associated weight *W*_*i*_ for each of the trees (usually set to 1), Min-Cut Supertree constructs the weighted proper cluster graph *G* = (*V, E, w*). The proper cluster graph contains vertices 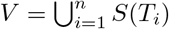. An edge (*u, v*) is in *E* if there exists any tree *T* ∈ 𝒯 for which *u* and *v* form a proper cluster. Let *I* be an indicator function which, given a tree and two taxa, returns one if the taxa form a proper cluster in the tree or zero otherwise. Each edge (*u, v*) ∈ *E* is weighted by the sum of the weights of the trees in which *u* and *v* for a proper cluster as demonstrated by Equation (1). An example proper cluster graph created from two trees is displayed in Figure 1.

**Figure 1:**
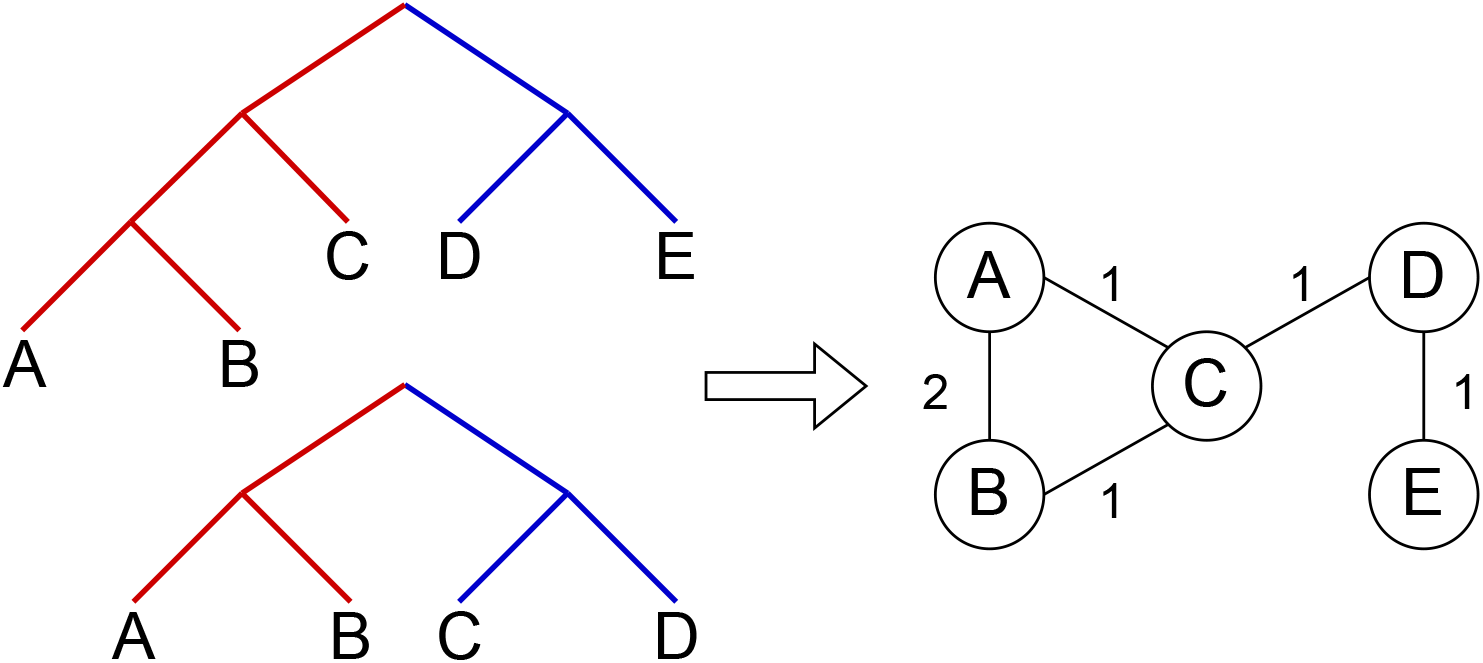
A proper cluster graph (right) for an example set of two source trees (left). The edges of the same colour identify taxa belonging to a proper cluster. Here, the trees have unit weights. That is, each edge in the proper cluster graph is weighted by the number of source trees in which the taxa form a proper cluster.

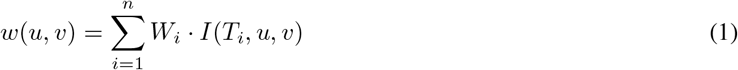

If there are any edges in the proper cluster graph (*e* ∈ *E*) with weight equal to the sum of the weights of the source trees 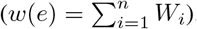, then every tree supports that proper cluster. Then, when finding a partition of the taxa, taxa in such proper clusters should never be separated. These edges are contracted to reduce the problem size. Figure 2 (left) shows the proper cluster graph from Figure 1 after such a contraction.

**Figure 2:**
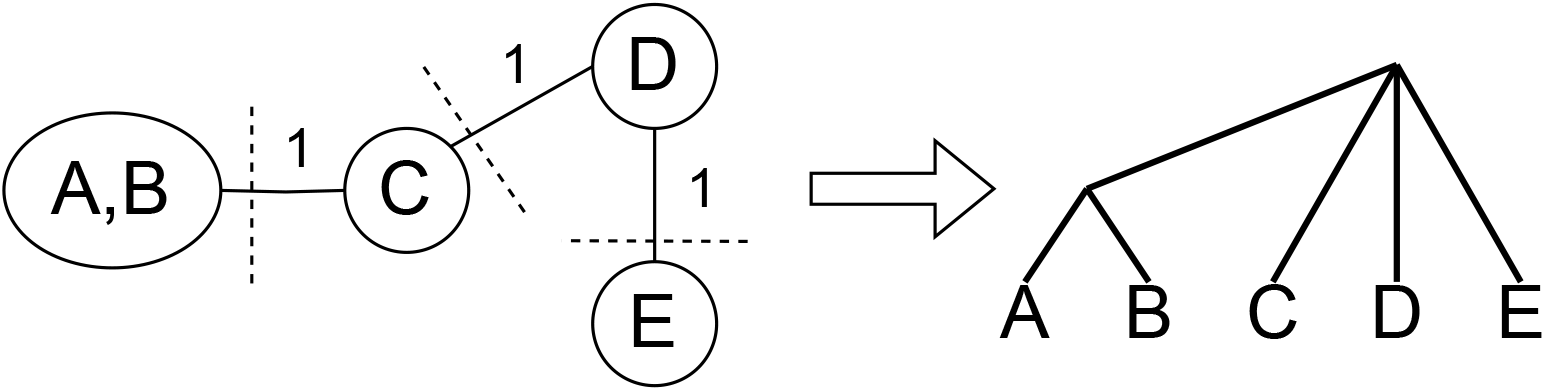
Graphical representation of the partitioning and result merging processes. After contraction, any edge on any min-cut of the proper cluster graph (left) is removed. Min-Cut Supertree is recursively called on the induced subtrees of the components of the proper cluster graph, making the roots of the resulting trees adjacent to a new root node (right).

#### 2.2.2 Finding the Best Partition

If the resulting proper cluster graph is disconnected, then the taxa in each of the components are always split over the roots of the source trees. Taxa in these components should thus be separated over the root of the supertree. Otherwise, there is no partition of the taxa for the root of the supertree satisfying the source trees. To resolve these cases, every edge that lies on any min-cut of the proper cluster graph is removed. This is analogous to removing groupings of taxa with the weakest support so that the graph can be partitioned.

The components of this graph partition the taxa into *n* disjoint sets {*X*_1_, *X*_2_, …, *X*_*n*_}. From this partition, *n* multisets of induced subtrees are generated {𝒯 |*X*_1_, 𝒯 |*X*_2_, 𝒯 |*X*_*n*_}. Min-Cut Supertree is then recursively called on each of these collections of induced subtrees.

If there are two or fewer taxa remaining when Min-Cut Supertree is called, the method immediately returns the phylogenetic tree containing those taxa. Once the trees generated from the recursive calls are returned, Min-Cut Supertree combines the results by connecting the roots of these trees to a new root node. This thus allows Min-Cut supertree to recursively create a complete rooted phylogenetic tree. Figure 2 demonstrates this, continuing the example in Figure 1.

### 2.3 Spectral Clustering

Spectral clustering [17, 27] is a clustering technique that can be applied to partition a graph over a “bottleneck”. We explain how spectral clustering works from two perspectives.

#### 2.3.1 Random Walk Perspective

We adapt this point of view from a tutorial on spectral clustering by Von Luxburg [33]. Consider an agent performing a random walk along a graph with *n* numbered nodes, and without loss of generality assume the graph is connected. The random walk can be represented as a symmetric stochastic matrix *P*. Let the vector *u* = (*u*_1_, *u*_2_, …, *u*_*n*_) represent the initial probability the agent is in each node. The probability the agent is in each node after *t* time steps is given by *P*^*t*^*u*. Spectral clustering can be used to partition the graph into two regions where the agent is most likely to remain trapped in for an extended number of time steps.

As *P* is a real symmetric matrix, it is eigendecomposable with eigenvalues *λ*_*i*_ and eigenvectors *v*_*i*_. Sort the eigenvectors by decreasing eigenvalue. By writing *u* in terms of *P* ‘s eigenvectors, the distribution of the agent after *t* time steps can be formulated as below.

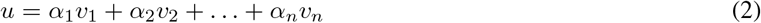

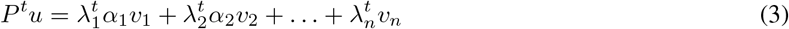

All eigenvalues of *P* are less than or equal to 1. The largest eigenvalue is 1, and the associated eigenvector is the stationary distribution *π* such that *Pπ* = *π*. For the symmetric stochastic matrix here, it is 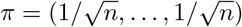. As *t* → ∞, the distribution of the agent converges to the stationary distribution. The second-largest eigenvalue/eigenvector pair thus represents the slowest part of Equation (3) to decay as it converges to the stationary distribution. The values in this eigenvector corresponds to “bottleneck” regions in the graph in which the agent may remain trapped in for the longest period of time steps. By partitioning this eigenvector into two clusters by, for example, using *k*-means clustering, the graph is separated over the bottleneck. This process can be applied in a similar fashion, starting from more general types of graphs [27]. It is usually done equivalently over the second-smallest eigenvalue of an alternative representation called the Laplacian matrix; see [14, 27, 33] for further reading.

#### 2.3.2 A Normalised Cut Perspective

Another perspective for looking at this problem is from the normalised cut perspective. Let *G* = (*V, E, w*) be a weighted graph. Define *W* : ℙ (*V*) *×* ℙ (*V*) → ℝ^+^ as the weighting between sets of vertices below:

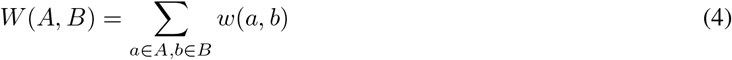

The normalised cut [27] of *G* aims to bipartition the vertices into two sets *A* and *B* such that the following formula is minimised:

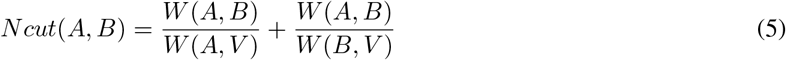

The solution to this optimisation problem essentially separates the vertices into the two most densely connected regions of the graph across a bottleneck. Minimising this value has been shown to be NP-complete [27]. However, a relaxation of the problem can be solved in polynomial time. The solution to the relaxation is the solution obtained through spectral clustering, yielding an efficient solution relying on eigensolvers [17, 33].

### 2.4 Proposed Algorithm: Spectral Cluster Supertree

Spectral Cluster Supertree is a supertree method derived from Min-Cut Supertree. Through analysis of the Min-Cut Supertree algorithm, it was determined that the most time-consuming operation was the min-cut step. In particular, the need to remove every edge on any min-cut of the proper cluster graph. The original paper [26] proposed a test for determining if an edge was in any min-cut of the proper cluster graph. For each edge, if the edge was deleted and the weight of the min-cut of the new graph was equal to the weight of the min-cut of the original graph minus the weight of the edge, then the edge is on a min-cut of the graph. Page [18] proposed an improvement to this method, using Picard and Queryanne’s algorithm [20] to explicitly find all minimum cuts of a graph. However, in practice, finding only a single arbitrary min-cut of the proper cluster graph in this algorithm was too computationally expensive.

Instead of relying on the (in practice) slow min-cut algorithm to partition the proper cluster graph, we use spectral clustering to efficiently separate the graph into two densely connected components. The remainder of the process is mostly identical to that of [26], using these components to induce the new trees for the recursive call. We also include an optimisation from Page’s modified Min-Cut Supertree [18] whereby if only one tree is present on a recursive call, it is returned early and grafted onto the growing tree. An overview of the algorithm is presented in Algorithm 1. We now present additional modifications that allow the method to make use of additional information provided by the source trees.

#### Algorithm 1

`SCS`(𝒯,W)

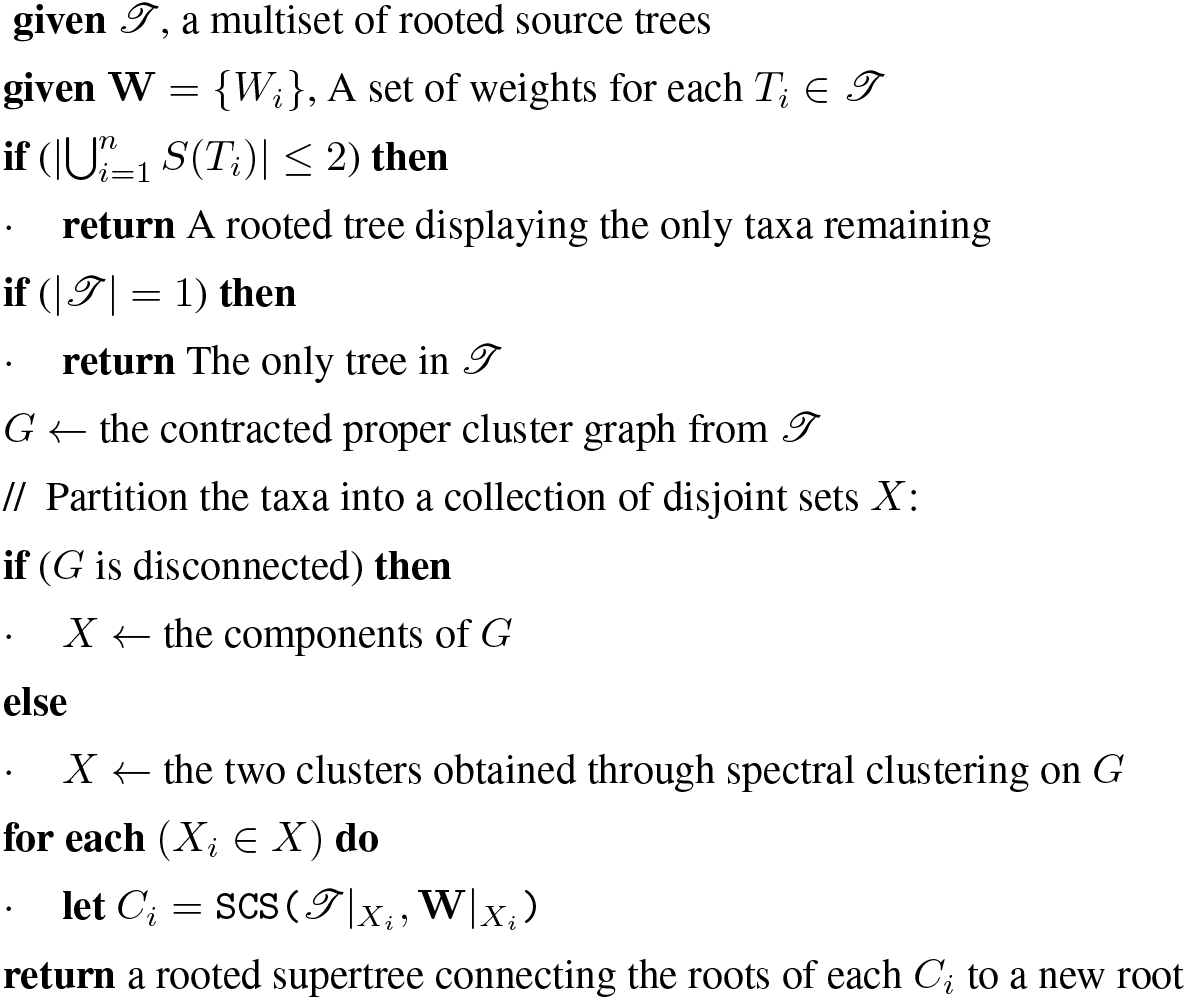

### 2.5 Weighting Strategies

BCD [7] introduced weighting strategies over the clades of the source trees to enhance the accuracy of their algorithm. Making use of the additional information that could be gleaned from the source trees allowed for more accurate supertree generation. To boost the topological performance of SCS, we outline similar weighting strategies gathered from information of the proper clusters in the source trees to weight the proper cluster graph.

For this part, recall that 𝒯 = {*T*_1_, *T*_2_, …, *T*_*n*_} is a multiset of rooted source trees, and *S*(𝒯) denote the set of taxa displayed in all of the trees. Like Min-Cut Supertree [26], we associate each source tree *T*_*i*_ ∈ 𝒯 with a user-specified weight *W*_*i*_. This can correspond to the user’s confidence in each of the source trees. If none are specified, unit weights are used. Finally, let lca be a function that, given a tree and two taxa, returns the node that is the lowest common ancestor of the two taxa if they are both present in the tree, or the root otherwise. We present two modifications to the weighting function for the proper cluster graph *w*(*u, v*), originally defined in Equation (1).

#### 2.5.1 Depth Weighting

SCS works by recursively splitting taxa over a root. The further the lowest common ancestor of a proper cluster is from the root of a source tree, the more likely that proper cluster is to be correct in the true tree. Conversely, the closer the lowest common ancestor of a proper cluster is to the root of a source tree, the weaker the proper cluster is. That is, it is more probable that the two taxa were incorrectly misplaced together on the same side of the root.

Depth weighting can be used when the source trees are provided without branch lengths, or when there is other reason for branch lengths to be ignored. Let *d* be a function that returns the depth of an internal node in a tree from its root (in terms of the number of edges from the root). When using depth weighting, the weight of each edge in the proper cluster graph for taxa *u* and *v* is given by Equation (6).

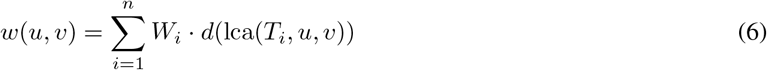

#### 2.5.2 Branch Length Weighting

Branch length weighting works under the same motivation as depth weighting, making use of the branch lengths of the source trees. Let *b* be a function that returns the branch length distance of an internal node in a given source tree from its root (the root node returns 0). The weight of each edge in the proper cluster graph for taxa *u* and *v* is given by Equation (7).

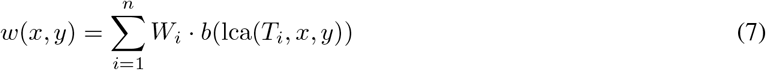

### 2.6 Implementation

We provide an implementation of our algorithm on GitHub under an open-source license. A Zenodo archive of the repository is available (https://doi.org/10.5281/zenodo.11118433). The algorithm was implemented using Python 3 and is installable as a package, sc-supertree, on PyPI. The Spectral Clustering step of the algorithm was implemented using scikit-learn [19]. The algorithm can be invoked through the command line using the scs command in its installed environment, taking as input a file name containing line-separated Newick-formatted source trees. The algorithm can also be utilised as a library with the `construct_supertree` function, taking as input a sequence of `cogent3` tree objects [12]. We also provide a `load_trees` function to convert a line separated Newick-formatted source tree file into a list of cogent3 trees. Further details on usage, including specifying the weighting strategy, are available in the project’s `README`.

## 3 Experimental Design

We perform extensive comparisons between SCS and BCD to evaluate their statistical and computational performance. Comparisons were made over a number of datasets, both existing and new. Our new dataset differs from those previously studied as they are generated without relying on an outgroup, and in that it aims to mimic what may be encountered during the last stage of a divide-and-conquer algorithm such as DACTAL [16]. This also corresponds to the practical application of inferring the tree root without prior knowledge of the phylogeny. All datasets are accessible online (https://doi.org/10.5281/zenodo.11118022).

### 3.1 Datasets

#### SMIDGenOG

The SMIDGenOG dataset [6] was developed using the SMIDGen protocol [30] in a rooted context using outgroup rooting. Datasets generated using the SMIDGen protocol aim to emulate data collection processes typically used by systematists, having many densely-sampled clade-based source trees, and a more widely sampled scaffold tree representing relations at a higher taxonomical level. Each model tree is paired with a single such scaffold tree and a number of densely sampled clade-based source trees. The scaffold tree samples a percentage of taxa, the scaffold-factor, uniformly at random over the entire tree. Universal genes (those that appear at the root and don’t become extinct) and non-universal genes were simulated along the model tree. These were used to generate alignments for the source trees. The trees were then constructed using RAxML [28]. All trees were rooted in this process using an outgroup, and the outgroup was subsequently removed. There are 30 model trees with corresponding source trees for every combination of 100, 500, and 1000 taxa with a scaffold factor of 20%, 50%, 75% and 100%.

#### SMIDGenOG-5500

The SMIDGenOG-5500 dataset [7] was created to be a large-scale dataset for the purposes of evaluating BCD. The dataset also aims to emulate approaches used by systematists but at a much larger scale. It was created using a similar methodology to SMIDGenOG. The dataset contains 10 model trees with an average of 5500 taxa. Each model tree is paired with 500 densely sampled clade-based source trees with 75-125 taxa. Due to the size of the dataset (to ensure the maximum likelihood process could run in a reasonable timeframe), each model tree is also paired with 5 sparsely-sampled scaffold source trees with 100 randomly selected taxa. This is instead of using one scaffold tree with a scaffold factor as in SMIDGenOG [6].

#### SuperTriplets

The SuperTriplets dataset [22] explores the effect of both the size of the source trees, and the number of source trees, on supertree construction. It contains 100 model trees with 101 taxa each (including an outgroup). The source trees are divided into a number of deletion rates *d* ∈ {25%, 50%, 75%} and a constant *k* ∈ {10, 20, 30, 40, 50}. In the data generation process, each model tree was duplicated 50 times with varied branch lengths. Sequence alignments were simulated on these duplicated trees. Maximum likelihood trees were generated from these 50 alignments, each under the constraint of *d* percent of the ingroup taxa being removed. The first *k* of these trees formed the set of source trees.

#### SCS Datasets

The SCS datasets aim to evaluate supertrees in a general setting where the data generation process does not necessarily rely on an outgroup. The datasets mimic what may be encountered by rooted variants of divide-and-conquer methods for phylogenetic reconstruction, such as DACTAL [16].

Ten model trees (with taxa 500, 1000, 2000, 5000, 10000) were generated following a birth-death process with a birth rate of 1.0 and a death rate of 0.2. The rooted ultrametric trees were initially scaled such that every tip was of distance one from the root. Similar to the SMIDGen protocol [30], branch lengths were then scaled by a random scaling factor. At the root, the scaling factor was 1.0 and the scaling factor evolved down different parts of the tree by adding a number from a normal distribution with a mean of 0 and a standard deviation of 0.05. The scaling factor was additionally bounded in this process between 0.05 and 8.0 to guard against excessively long or short branch lengths following the SMIDGen protocol.

For each of the model trees, a sequence alignment of length 10000 was simulated using a strand-symmetric general nucleotide Markov substitution process under which the root is identifiable [10]. The alignment length was chosen to allow for a sufficient level of accuracy when estimating the source trees. The parameters for the process were estimated from a sequence alignment of 3 bacterial species [11].

Rec-I-DCM3 [24] is a method that can be used to decompose a tree into overlapping subsets of taxa. We have made an implementation of this algorithm in Python publicly available (https://doi.org/10.5281/zenodo.11118313). We used Rec-I-DCM3 to partition each model tree into overlapping subsets of taxa of maximal sizes 50 and 100. We extracted the subtrees for these partitions from the model tree to form a collection of source trees. This forms a simple dataset we call the “SCS-Exact” dataset, from which all methods should always be able to reconstruct the true tree.

For the more interesting second dataset, for each subset in the partition of taxa formed by Rec-I-DCM3, the corresponding simulated sequences were extracted. IQ-TREE 2 [15], under a strand symmetric model, was used to fit a rooted tree to these sequences. The set of trees generated for a partition formed the source trees for the “SCS-DCM-IQ” dataset.

### 3.2 Distance Measures

We measure the topological accuracy of supertrees generated by the evaluated methods using the standard Robinson-Foulds distance [23], but also the more statistically robust Matching Cluster distance [3]. For this paper, we use a definition of Robinson-Foulds that is the cardinality of the set containing the symmetric difference of the clades between the two compared trees. Let *C* be a function mapping trees to their set of clades, and ⊕ denote the symmetric difference operator. The Robinson-Foulds distance is given by Equation (8).

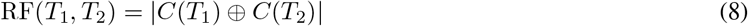

The Robinson-Foulds distance, while quick to compute, is known to exhibit poor statistical behaviours [2, 13]. For instance, it is known to saturate quickly. This means similar topologies differing by only a single leaf can maximise this metric, as illustrated by Figure 3.

**Figure 3:**
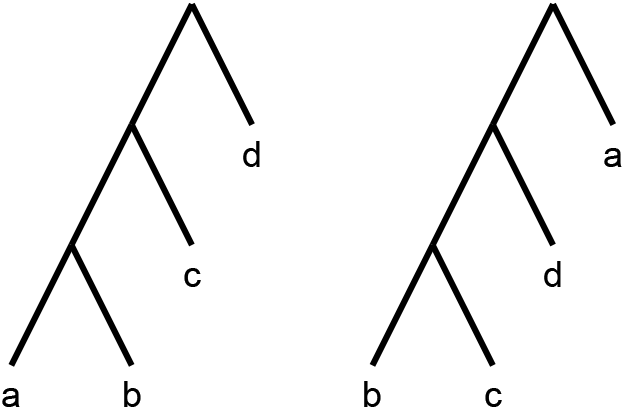
By moving a single leaf (a) to the root of the tree, the Robinson-Foulds distance is maximised.

Consider the F_1_ score used by Fleischauer and Böcker [7], and adapt it to the rooted case (the unrooted version and unrooted Robinson-Foulds follows similarly). This adaptation can be done by letting “true positives” (TP) refer to clades in both the supertree and model tree; “false positives” (FP) refer to clades in the supertree but not the model tree; and “false negatives” (FN) refer to clades in the model tree but not the supertree. Then, the Robinson-Foulds distance is simply RF = FP + FN. Further, there is a direct mapping between the Robinson-Foulds distance and F_1_ score scaled by the number of correctly identified clades.

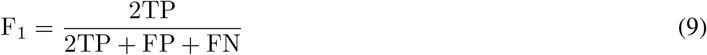

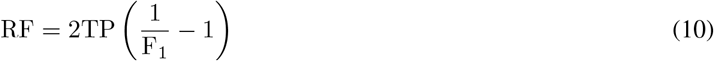

Further, the number of non-root internal nodes in the supertree is equal to the sum of the number of true positive and false positive clades. In the case of a fully resolved tree with *n* taxa containing no polytomies, there are *n* − 2 such internal nodes and 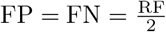. Thus, 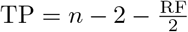 and, treating the number of taxa as constant, there is an exact linear mapping between the F_1_ score and Robinson-Foulds distance given by Equation (11). This implies that under this scenario, the F_1_ score and Robinson-Foulds distance are, in effect, measuring the same relation. Though this exact mapping assumes no polytomies, in practice when we calculated the Robinson-Foulds distance and F_1_ score they told very similar stories in the results. As such, we only report the Robinson-Foulds distance of these two in our comparisons and include the F_1_ score in the supplementary material.

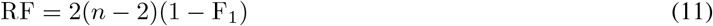

We will also include the strictly more statistically robust Matching Cluster distance in our comparisons. The Matching Cluster distance [3] is similar to the *β* distance [4]. Rather than considering only exact matches of clades, it also takes into consideration the degree of dissimilarity between the clades. It does this by solving a min-weight matching problem between the symmetric differences of each pair of clades in the two trees. For the example in Figure 3, the non-trivial clusters are {{*a, b*}, {*a, b, c*}} and {{*b, c*}, {*b, c, d*}} respectively. Figure 4 shows a bipartite graph between the non-trivial clusters of the two trees. The edges are weighted by the cardinality of the symmetric difference of the clusters. The solution to the min-weight matching problem here is 4. The maximum possible Matching Cluster distance for bifurcating trees with 4 taxa here is 6.

**Figure 4:**
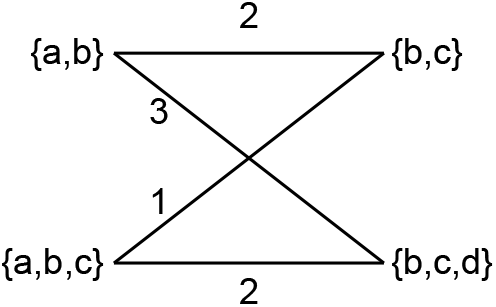
The Matching Cluster distance is calculated by solving the min-weight matching problem of a bipartite graph. Each side of the bipartite graph contains the clusters of the respective trees. Edges are weighted by the cardinality of the symmetric distance of the sets it connects.

When considering multifurcating trees, one tree may have fewer clades than the other. This could mean one side of the graph in the min-weight matching problem could have fewer nodes than the other. The Matching Cluster distance generalises to such scenarios by adding empty sets to the side with fewer nodes until the number of nodes on each side are equal.

The Matching Cluster distance is strictly more robust than the Robinson-Foulds distance. In fact, for bifurcating trees, if the weights of the edges for clusters are replaced by 0 if they do match, and 2 if they do not, the Robinson-Foulds distance and Matching Cluster distance become identical.

### 3.3 Experiments

Experiments were performed using the NCI’s Gadi (https://nci.org.au), running on a single core of an Intel(R) Xeon(R) Platinum 8268 CPU @ 2.90GHz with 16 GB of RAM. SCS, BCD with and without the Greedy Strict Consensus Merger (GSCM) preprocessing, and Min-Cut Supertree (MCS) were evaluated over all of the datasets. In particular, a modified version of Min-Cut Supertree was used that included our weighting strategies for the proper cluster graph. We also allowed MCS to find only a single arbitrary min-cut of the proper cluster graph, rather than all min-cuts, so the problems could be solved within a reasonable time. Branch length weighting was utilised across all methods where possible (all but the SuperTriplets dataset). Otherwise, depth weighting for our method, or unit weighting for BCD, was used. The CPU time to resolve each supertree was recorded, as well as the Robinson-Foulds Distance and Matching Cluster Distance when compared to the model tree. (F_1_ score results are included in the supplementary material). If a method did not complete a dataset under specific parameters within a wall time of 48 hours, it was terminated early. We have made the code used to process and evaluate the experiments is publicly available (https://doi.org/10.5281/zenodo.11118313). It also includes a script for downloading the datasets.

## 4 Results

### 4.1 Spectral Cluster Supertree vs Min-Cut Supertree

Here, we show how SCS is a significant improvement over Min-Cut Supertree across all measures. Figure 5 displays the difference in the recorded measures between the two methods over the largest dataset where Min-Cut Supertree resolved at least some problems (4/10 for each of the maximum subproblem sizes). Where SCS takes a matter of seconds to solve a problem, Min-Cut Supertree can take from two hours to close to a full day. SCS additionally outperforms Min-Cut Supertree in every metric – consistently producing more correct clades in the generated supertree (from the Robinson-Foulds Distance), and markedly superior general topological accuracy from the Matching Cluster Distance.

**Figure 5:**
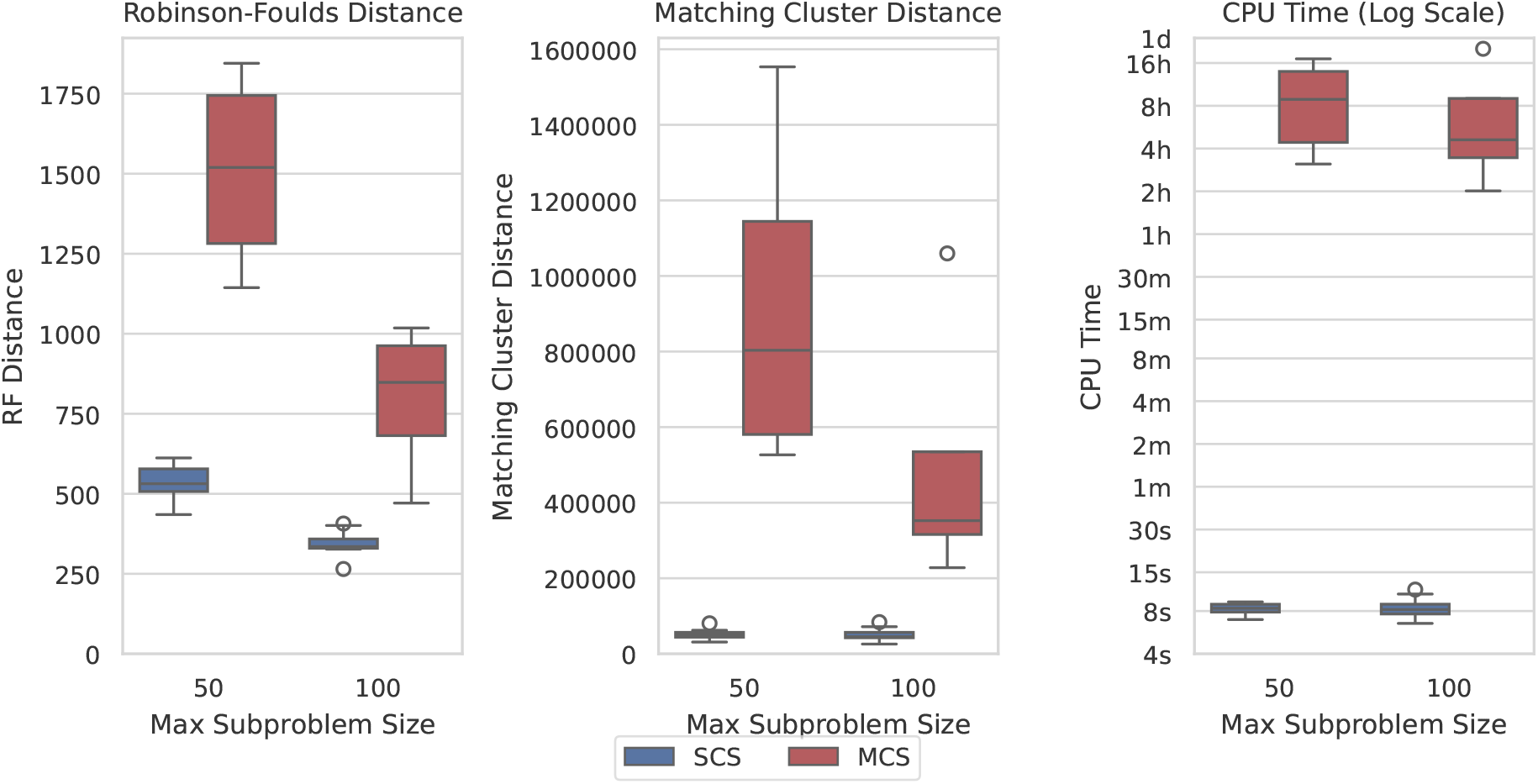
SCS outperforms MCS on the SCS-DCM-IQ dataset with 5000 taxa. Lower is better for all graphs. Time results are shown on a log scale. While SCS solved all problems, MCS only solved 4/10 within the time limit for each of the maximum subproblem sizes.

The results shown in Figure 5 are similar across most datasets, though not necessarily to the same extent with regard to time on the smaller datasets. MCS only outperformed SCS in time in the SCS-Exact dataset, where importantly no min-cut or spectral clustering calls are required, and all methods recovered the true tree. On the largest SCS-Exact problems, both methods took around 6-10 seconds. This illustrates the extent at which the min-cut component of the MCS impacts its computational efficiency. SCS always dominated MCS with respect to topological accuracy under both distance metrics on all other datasets. Going forward, we thus only compare the performance of SCS to BCD.

### 4.2 Spectral Cluster Supertree vs Bad Clade Deletion

All methods could construct the correct supertree in the SCS-Exact dataset, though GSCM preprocessing made BCD sometimes take hours for the largest problems. Without GSCM, BCD finished in a few seconds. We now show the results for SCS compared with BCD with and without GSCM preprocessing over each of the datasets.

Figure 6 shows the results over the SCS-DCM-IQ datasets with 10000 taxa. Results for the other taxa counts follow the same pattern (see figures A.1-A.4). The figure shows that with GSCM preprocessing, BCD outperforms SCS in terms of Robinson-Foulds distance. This metric measures the number of clades that are different (no matter how similar) between the model tree and generated supertree. SCS however outperforms BCD under the Matching Cluster distance, which measures more generally how similar the overall topologies of the tree are - taking into account the degree of similarity/dissimilarity in the clades. SCS is also vastly superior here compared to BCD in terms of the time taken to solve the problems. Where BCD takes multiple hours per problem instance, SCS can solve these problems in less than a minute - usually in under twenty seconds.

**Figure 6:**
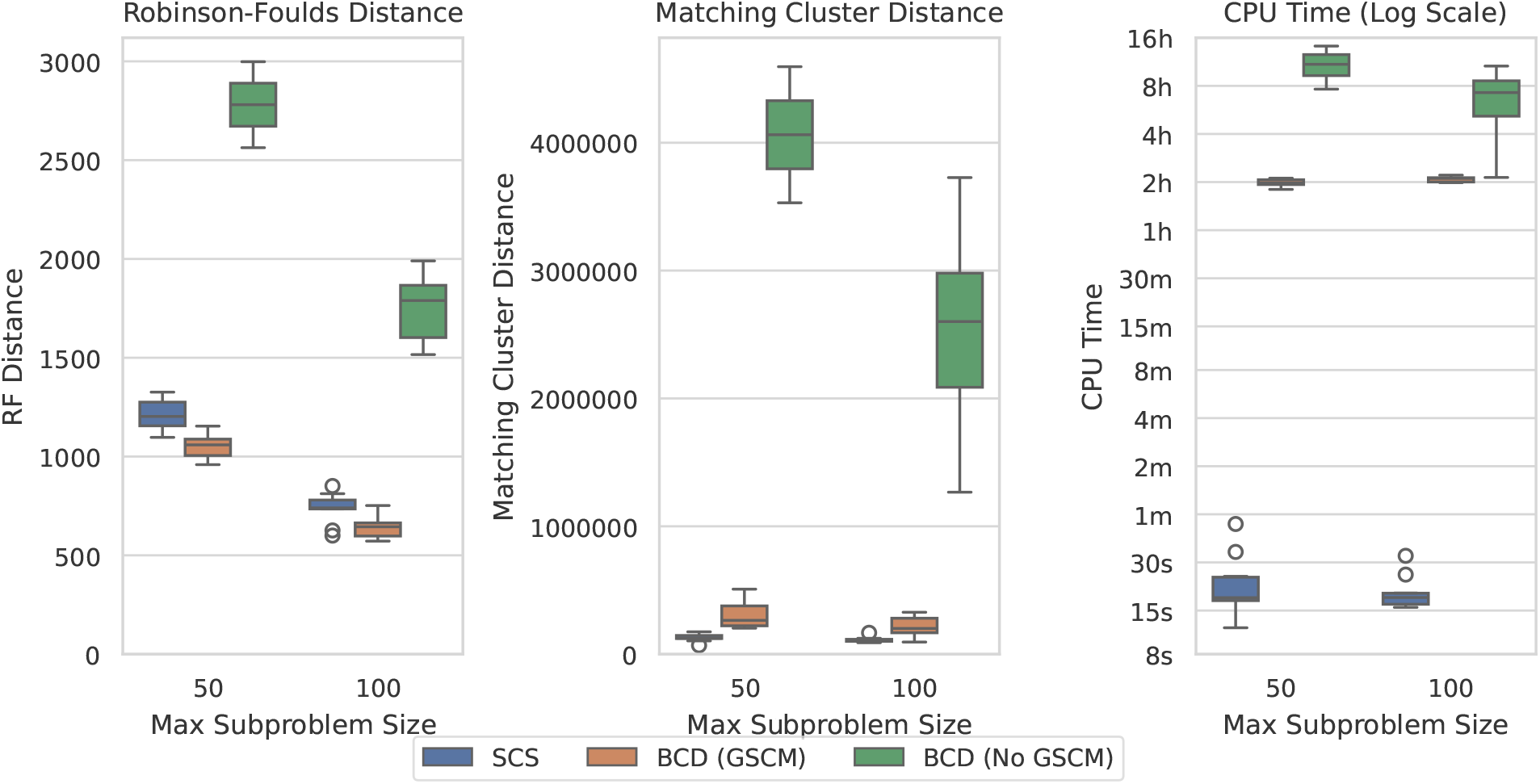
SCS vs BCD on the SCS-DCM-IQ dataset with 10000 taxa. The dataset emulates what may be encountered when using divide-and-conquer algorithms for phylogenetic reconstruction. BCD without GSCM processing only solved the first 2 and 6 problems within the timeout for the 50 and 100 max subproblem sizes respectively. The other methods solved all ten problems.

Figure 7 compares the results of SCS to BCD under the large SMIDGenOG-5500 dataset. The results follow a similar pattern to the SCS-DCM-IQ dataset. BCD still outperforms SCS in terms of the Robinson-Foulds Distance, though the difference is less pronounced. SCS still outperform BCD in terms of the Matching Cluster distance, which compares the tree topologies more generally. SCS is still a vast improvement over BCD in terms of the time required to solve the problems, taking minutes, where the others can take multiple hours.

**Figure 7:**
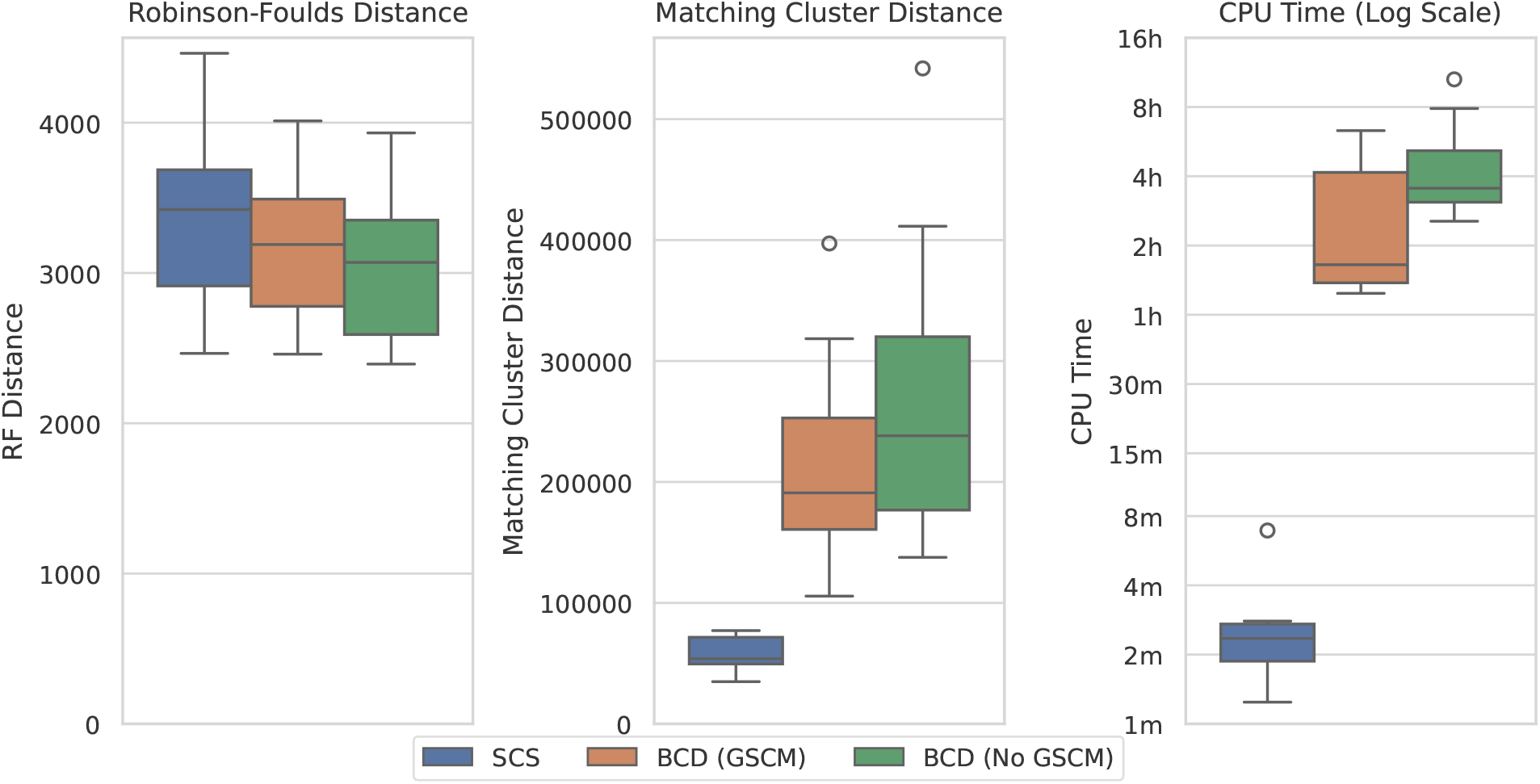
SCS vs BCD on the SMIDGenOG-5500 dataset. The dataset applies the SMIDGen protocol to mimic what may be encountered by systematists at a large scale. As there is no parameterisation for this dataset, no x-axis is displayed. All methods solved all problem instances within the timeout.

Figure 8 shows the results over the largest original SMIDGenOG dataset with 1000 taxa and different scaffold factors (the percentage of taxa sampled for the scaffold trees). This dataset shows the best results for BCD when compared with SCS. BCD outperforms SCS in terms of the Robinson-Foulds Distance. The results are roughly even in terms of Matching Cluster Distance for the smaller two scaffold factors, though BCD with GSCM preprocessing is slightly ahead with the higher two scaffold factors. SCS is slightly ahead on time compared with BCD with GSCM preprocessing. However, problems here are being solved in the order of seconds and the difference is not significant for practical purposes. Results are similar for the 100 and 500 taxa counts (see appendix figures A.5 and A.6), with BCD being faster on the 100 taxa dataset.

**Figure 8:**
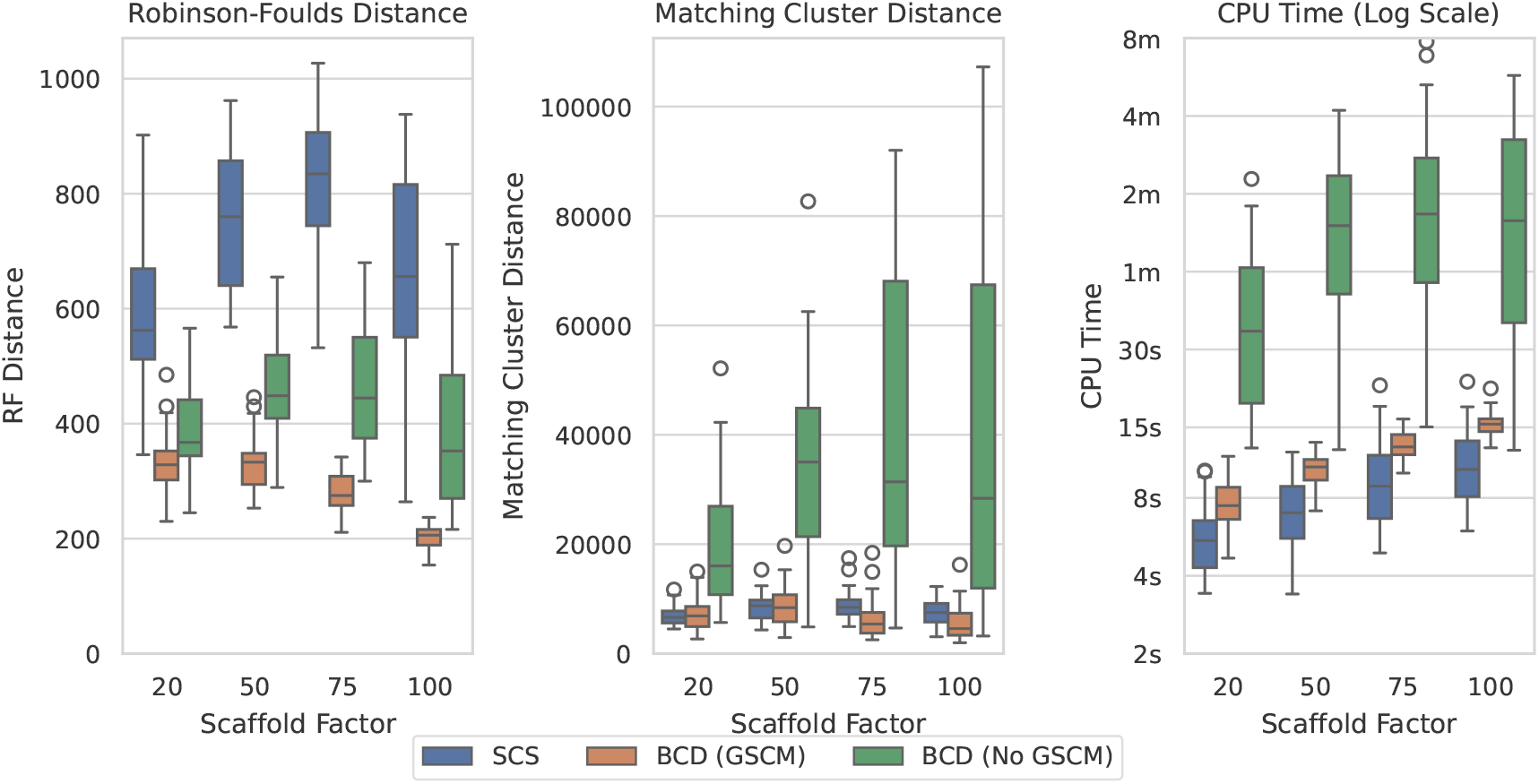
SCS vs BCD on the SMIDGenOG dataset with 1000 taxa. The dataset aims to imitate data curation processes of systematists. The SMIDGenOG dataset contains one scaffold tree sampling over “Scaffold Factor” percent of the taxa, as well as many densely sampled clade-based source trees.

Figure 9 shows the results over the SuperTriplets dataset with a 50% deletion rate (percentage of taxa removed when generating each source tree). The SuperTriplets dataset shows a significant amount of variance over the different deletion rates. With a 50% deletion rate, SCS generally performs worse with respect to the Robinson-Foulds Distance, but better in terms of Matching Cluster distance. With a 75% deletion rate (appendix figure A.8), SCS generally outperforms BCD in all distance metrics. For the 25% deletion rate (appendix figure A.7), BCD outperforms SCS in terms of the Robinson-Foulds distance. The central distribution of the Matching Cluster distance is roughly identical, with BCD achieving better minimum values, though SCS achieves better maximum values. The time results here are all low enough to not make much of a difference for practical purposes.

**Figure 9:**
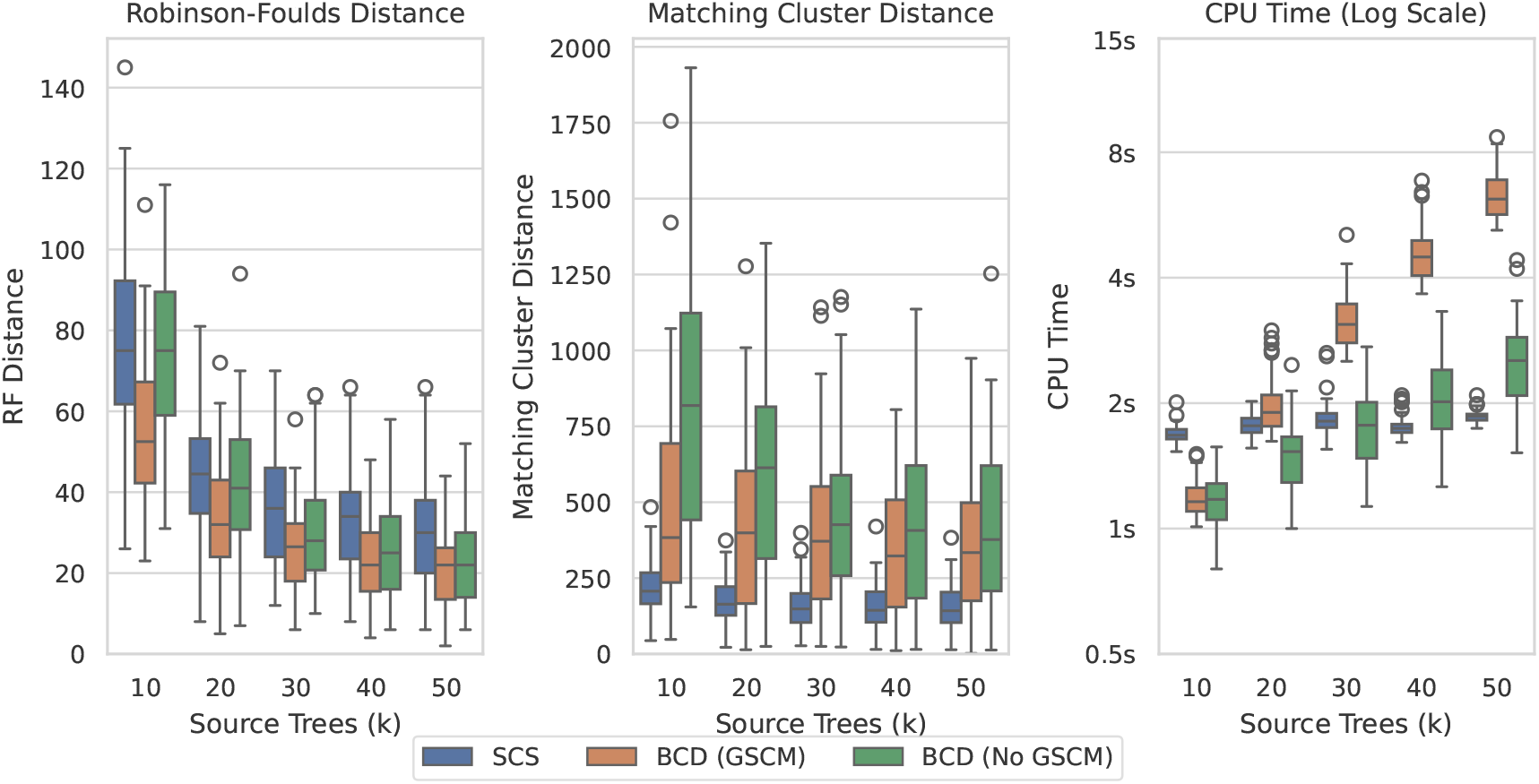
SCS vs BCD on the SuperTriplets dataset with a deletion rate of 50%. The SuperTriplets dataset explores the effect of percentage of taxa in the source trees, and number of source trees, on supertree construction. Results are variable dependent on the deletion rate, and results for the others are included in the appendix.

Full figures for the other parameterisations of the datasets are available in the appendix. The raw results have been archived on Zenodo (https://doi.org/10.5281/zenodo.11118313).

## 5 Discussion

The Spectral Cluster Supertree (SCS) algorithm for merging rooted phylogenetic trees exhibits comparable or markedly better statistical and computational performance than current approaches. While Min-Cut Supertree (MCS) was demonstrated impractical for modern sized data sets, Bad Clade Deletion (BCD) was far more efficient when used in conjunction with its GSCM pre-processing step. Under most conditions examined, particularly for large problem size, SCS was orders of magnitude faster than BC. Comparison of the statistical performance of the algorithms was sensitive to the topology distance metric used. Thus, judgement of the statistical merits of SCS relative to competing approaches hinges on the properties of the Robinson-Foulds and Matching Cluster distance metrics.

### SCS Statistically Outperforms BCD on Most Datasets

In terms of Robinson-Foulds distance, BCD with GSCM preprocessing almost always outperformed SCS or were otherwise roughly equivalent. However, the Robinson-Foulds distance metric suffers from poor statistical qualities. It can saturate quickly, it is possible for trees that are identical minus the placement of a single leaf to maximise this distance metric. It simply compares how many of the clades are an exact match between the model tree and supertree, without considering the degree of similarity or dissimilarity between the clades.

The Matching Cluster distance is a strictly more robust distance measure. It gathers the clades of the model tree and supertree, and measures the degree of dissimilarity between the clades of these two sets (based on the size symmetric difference of the clades, or in other words, how many taxa in the clades are different). By then finding the optimal pairing of clades between these sets, the distance measure is calculated. By measuring the degree of similarity between the clades of the trees, it is a strictly superior measure. If instead the degree of dissimilarity was measured as 2 if the clades do not match and 0 otherwise, the distance measure would become identical to Robinson-Foulds.

SCS frequently outperformed BCD in terms of Matching Cluster distance on all datasets but the SMIDGenOG dataset, and Supertriplets dataset with a deletion rate of 25%. When measuring the median improvement of this metric over each parameterisation of the datasets on a problem by problem basis, SCS was 1.10-2.58 times better on the SCS-DCM-IQ dataset; 3.45 times better than BCD on the SMIDGenOG-5500 dataset; and anywhere from 1.93-2.25 times and 2.60-6.60 times better on the Supertriplets dataset with a deletion rate of 50% and 75% respectively. On the other datasets, BCD mostly performed better than SCS by 0.97-1.68 times on the SMIDGenOG dataset, and 1.18-1.43 times on the Supertriplets dataset with a deletion rate of 25%.

It is somewhat curious, on first consideration, the discrepancy between the SMIDGenOG and SMIDGenOG-5500 results given the underlying data was simulated under the same protocol. The primary difference between the two datasets here is the number of taxa in the scaffold tree. Due to practical limitations, SMIDGenOG-5500 contains 5 scaffold trees with 100 taxa each (less than 2% of the taxa each on average) and SMIDGenOG contains a single scaffold tree containing either 25%, 50%, 75% or 100% of the taxa. It also appears the rate at which BCD improves on SCS decreases as the scaffold factor decreases. Further, the effect of the deletion rate under the Supertriplets dataset indicates that the more taxa that are removed from each of the source trees (higher deletion rate), the more topologically accurate SCS is compared to BCD.

The algorithmic properties of the two methods could potentially explain this relationship. BCD effectively aims to delete a minimum number of clades from the source trees when conflicts arise to construct a supertree. Importantly, every clade in the supertree must also be a clade in one of the source trees (which leads to lower in comparison Robinson-Foulds distances). Having a strong backbone with a widely sampled scaffold tree can support this. When conflicts arise in SCS, however, we consider the degree to which the source trees support taxa being grouped on the same side of the rooted tree as other taxa. We partition the taxa, into two groups which maximal support within each group but minimal support between the two groups. This is clearly quite effective in practice, but it may not count as heavily, in comparison, the presence of a strong backbone. On further examination. we found that by increasing the tree weight associated with the scaffold tree in the SMIDGenOG dataset, the accuracy of the SCS method improved significantly (sometimes better than BCD). The improvement was particularly strong for high scaffold factors. This supports our hypothesis for the explanation of the relationship here. However, as BCD also supports tree weights and to avoid overfitting these extra parameters to the reported accuracy, we do not include these results in our comparisons.

When considering practical applications of the algorithms on large datasets, the choice of method is clear. For large phylogenetic reconstruction problems, the time required to perform a full maximum likelihood analysis over sequences of interest is far too computationally taxing. In these scenarios (as in the SMIDGenOG-5500 dataset), the presence of a large scaffold tree can potentially be infeasible. Divide-and-conquer methods relying on the Disk-Covering Method [24], breaking down the problem into small overlapping sub-problems, may thus be required to make computation possible. In these situations, it is clear SCS performs best with respect to topological accuracy, though BCD may still be useful for smaller problems with a strong scaffold tree.

### SCS Computationally Outperforms BCD

SCS is also vastly more efficient in terms of CPU Time compared to BCD. For the largest problems in the SCS-DCM-IQ dataset, SCS could solve problems that took BCD roughly 2 hours in under a minute - usually in less than 20 seconds (median speedup 386 and 409 for a 50 and 100 maximum subproblem size respectively). For the SMIDGenOG-5500 dataset, where BCD took 1-8 hours to solve each problem, SCS took only 1-8 minutes (median 58 times speedup). Ignoring problems which took both BCD (GSCM) and SCS under fifteen seconds to solve, the median speedup over BCD under each combination of parameters of the datasets ranged from 12-409 - the greatest speedups on the largest datasets. The only problems BCD beat SCS on speed, SCS still solved in only 1-2 seconds. SCS is clearly the superior choice in terms of time needed to fully resolve a supertree.

It must be noted that the results illustrated in the figures were obtained from running the experiments on a single CPU core. BCD has support for multi-threading, which is performed over both min-cut computations and recursive calls. In the current version of SCS, concurrency is only utilised within NumPy operations [8] and during a part of the spectral clustering step as implemented in scikit-learn [19]. There is room for further parallelisation in SCS, however. Similar to how BCD parallelises across independent recursive calls, the same can be done trivially for SCS. This parallelisation was trialled, but the improvement was minor on these problem sizes due to the associate overhead, and the time domination of performing the first and largest split. Parallelisation will likely have a much more beneficial effect for even larger problems than we have tested, and we accordingly leave a full comparison of the parallel versions for future work. Due to BCD’s parallelisation over solving min-cuts in addition to subproblems, greater speedup can be obtained, particularly on these problem sizes. However, SCS improves on the time results of BCD over large problems to such a significant extent that this difference may not matter. This is especially true when considering limitations such as Amdahl’s law (which gives a theoretical asymptotic limit to the speedup of a program as the number of processors are increased).

## 6 Conclusion

We presented a new algorithm, Spectral Cluster Supertree, for merging overlapping rooted phylogenetic trees. Our algorithm is significantly faster than Bad Clade Deletion on large problem sizes, taking on average 20 seconds where Bad Clade Deletion took ∼ 2 hours on one dataset, and 1-8 minutes where Bad Clade Deletion took 1-8 hours on another. While Bad Clade Deletion can sometimes display a superior topological accuracy on datasets containing large scaffold trees, on most datasets Spectral Cluster Supertree is more accurate. Of particular importance, Spectral Cluster Supertree was more topologically accurate than Bad Clade Deletion on large problems or otherwise those where the number of taxa in each of the source trees may be low in proportion to the total number of taxa. This may be especially valuable for large problems where divide-and-conquer methods for phylogenetic reconstruction could be necessary for computational feasibility. We leave further comparisons with respect to parallel implementations, where larger datasets than those currently considered are necessary for proper investigation.

## A Full SCS vs BCD Results

**Figure A.1:**
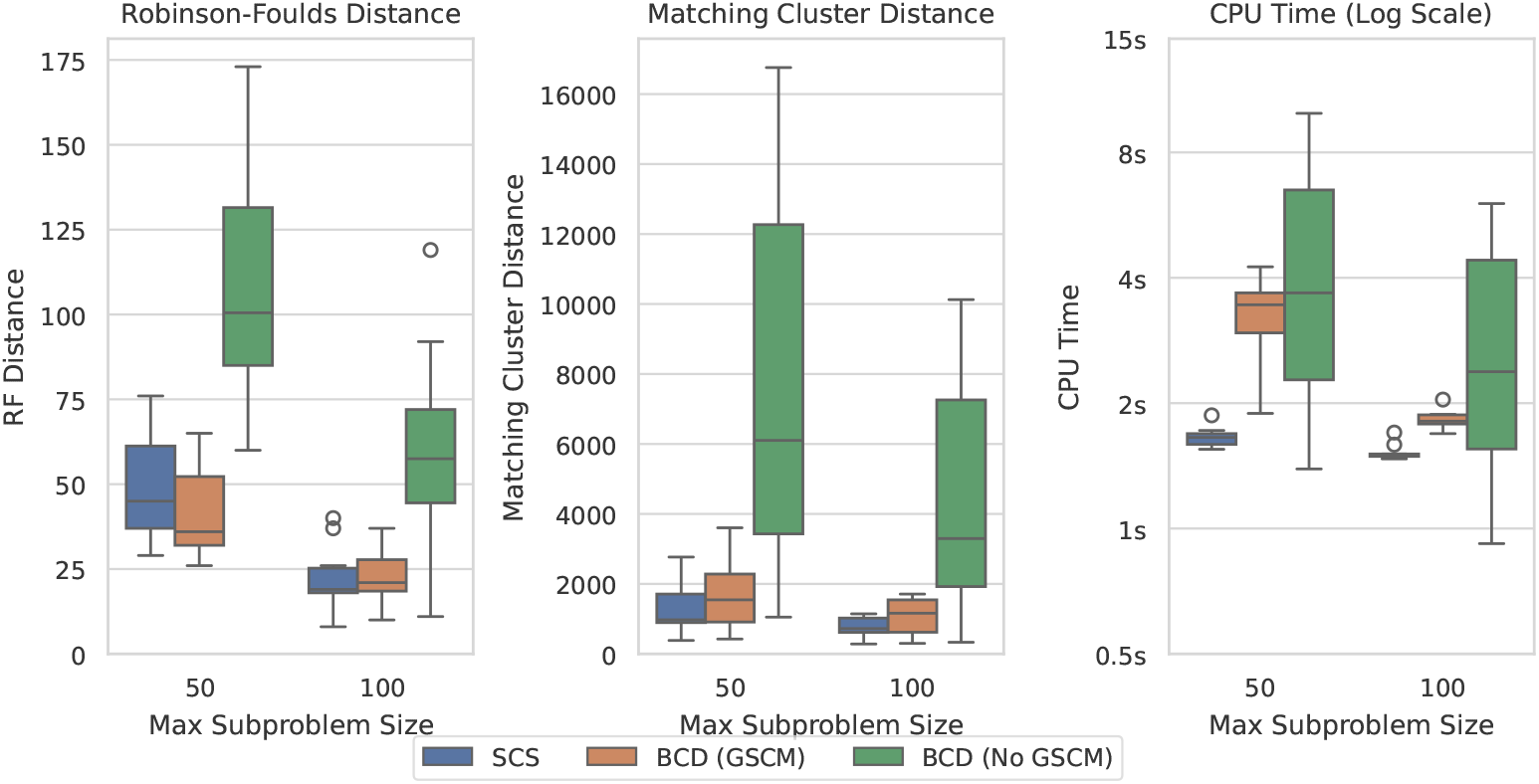
Spectral Cluster Supertree vs Bad Clade Deletion on the SCS DCM IQ-TREE dataset with 500 taxa.

**Figure A.2:**
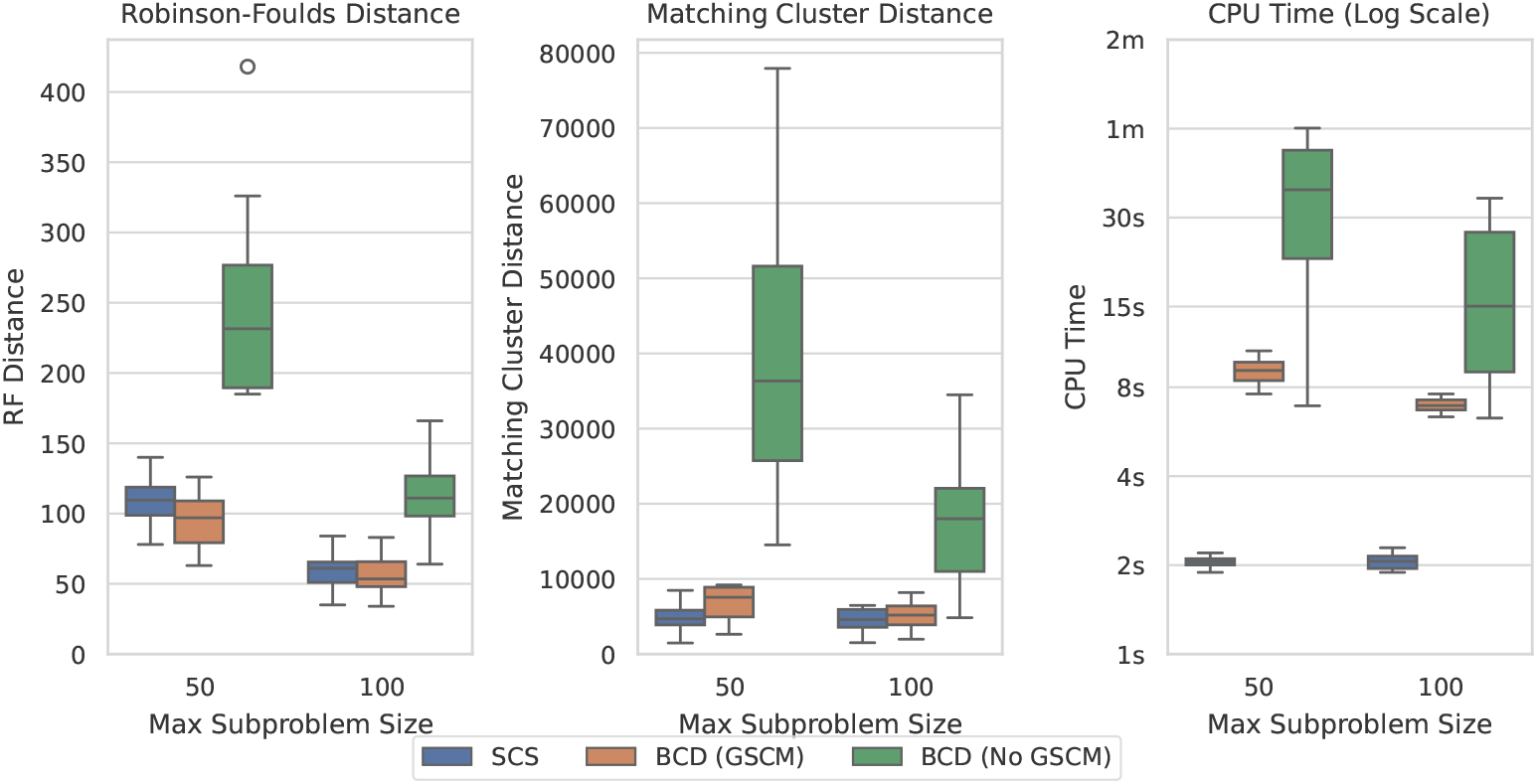
Spectral Cluster Supertree vs Bad Clade Deletion on the SCS DCM IQ-TREE dataset with 1000 taxa.

**Figure A.3:**
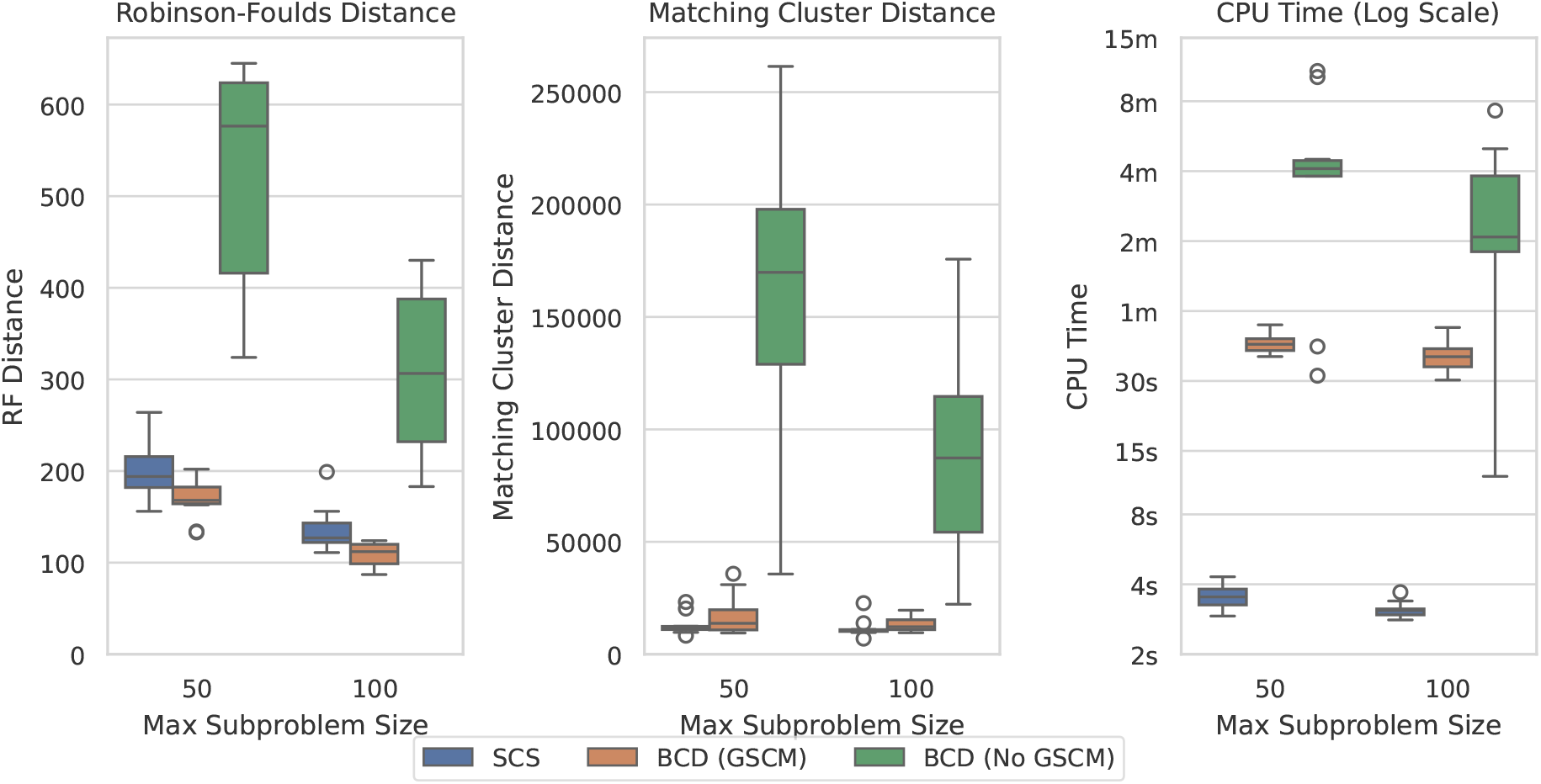
Spectral Cluster Supertree vs Bad Clade Deletion on the SCS DCM IQ-TREE dataset with 2000 taxa.

**Figure A.4:**
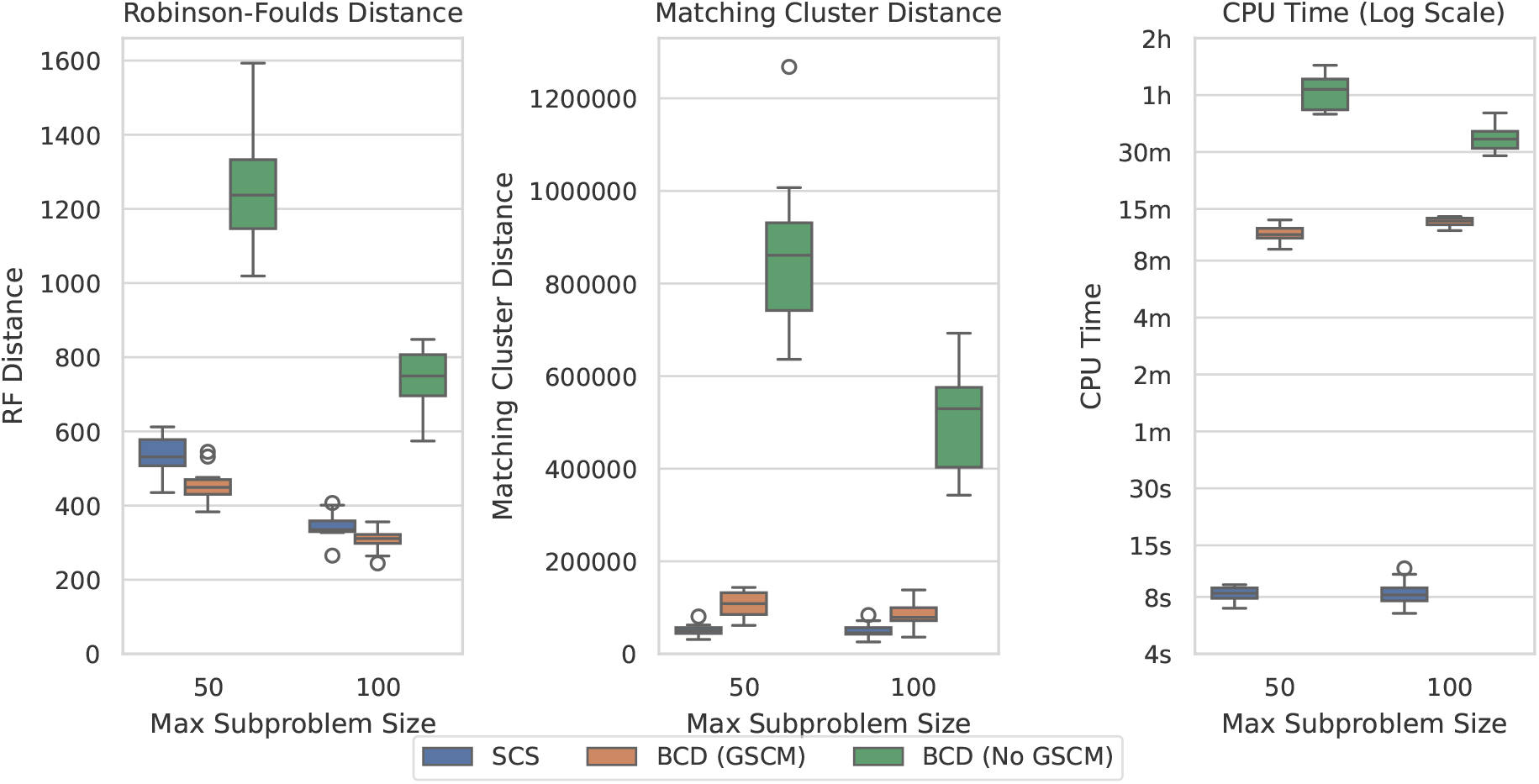
Spectral Cluster Supertree vs Bad Clade Deletion on the SCS DCM IQ-TREE dataset with 5000 taxa.

**Figure A.5:**
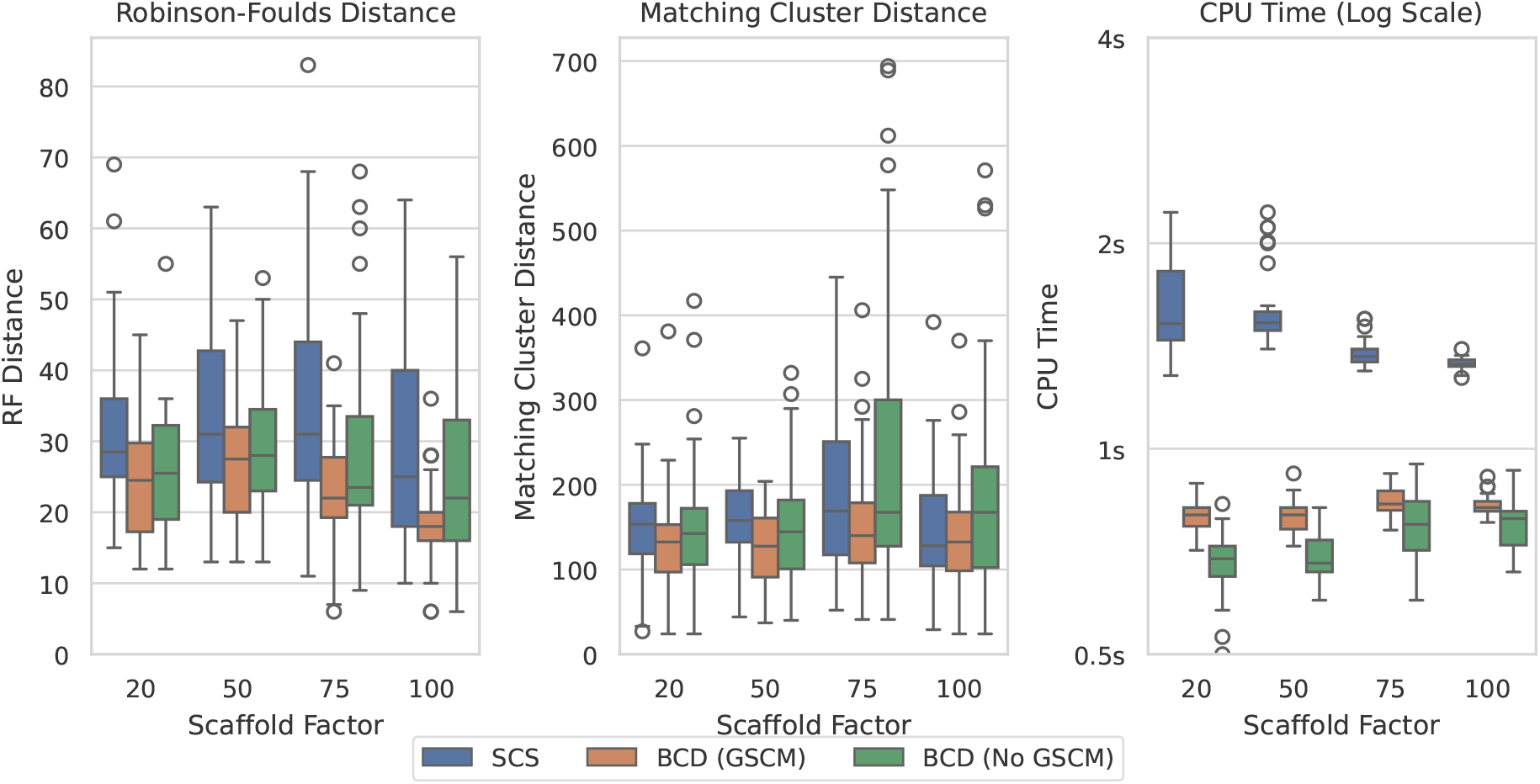
Spectral Cluster Supertree vs Bad Clade Deletion on the SMIDGenOG dataset with 100 taxa.

**Figure A.6:**
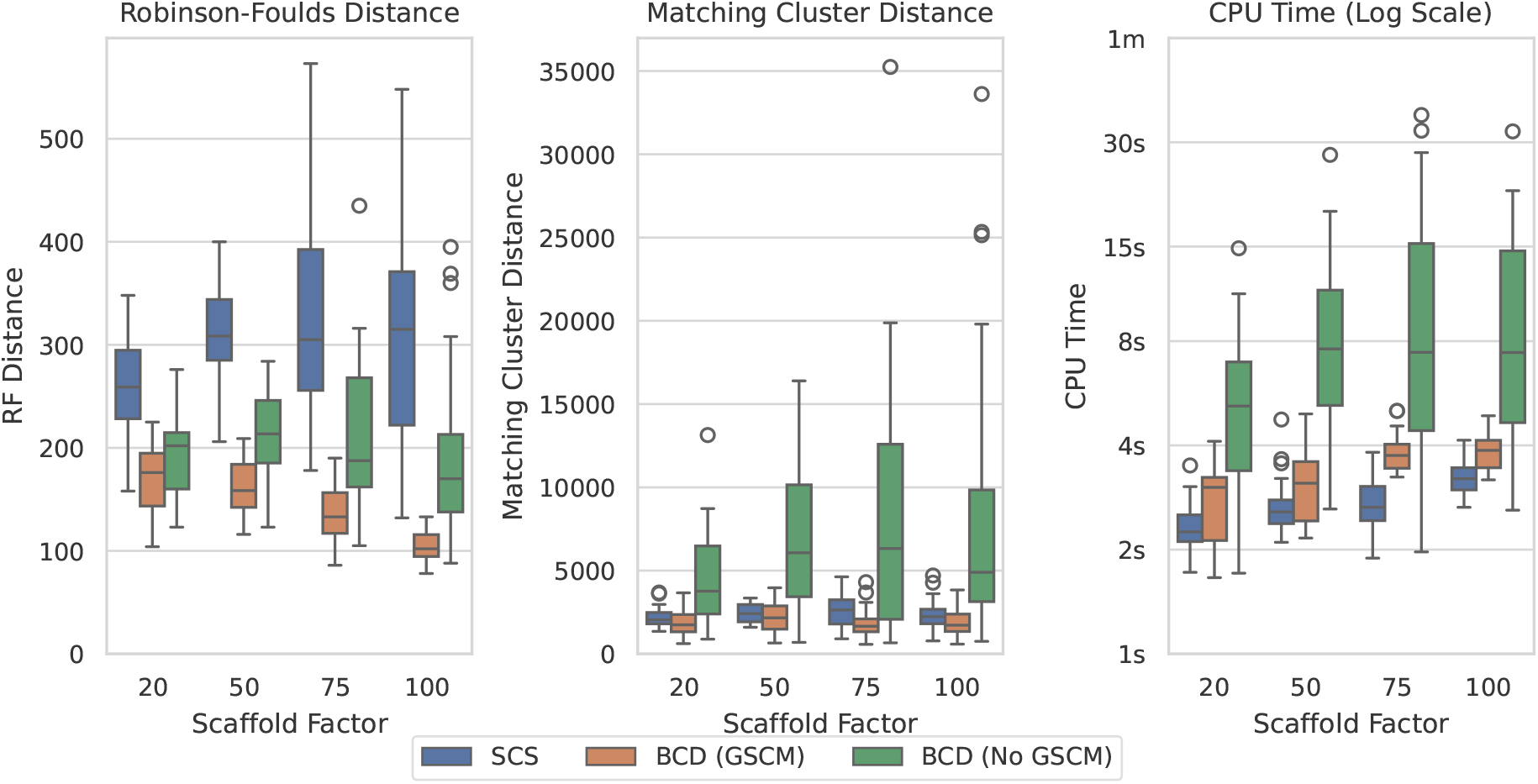
Spectral Cluster Supertree vs Bad Clade Deletion on the SMIDGenOG dataset with 500 taxa.

**Figure A.7:**
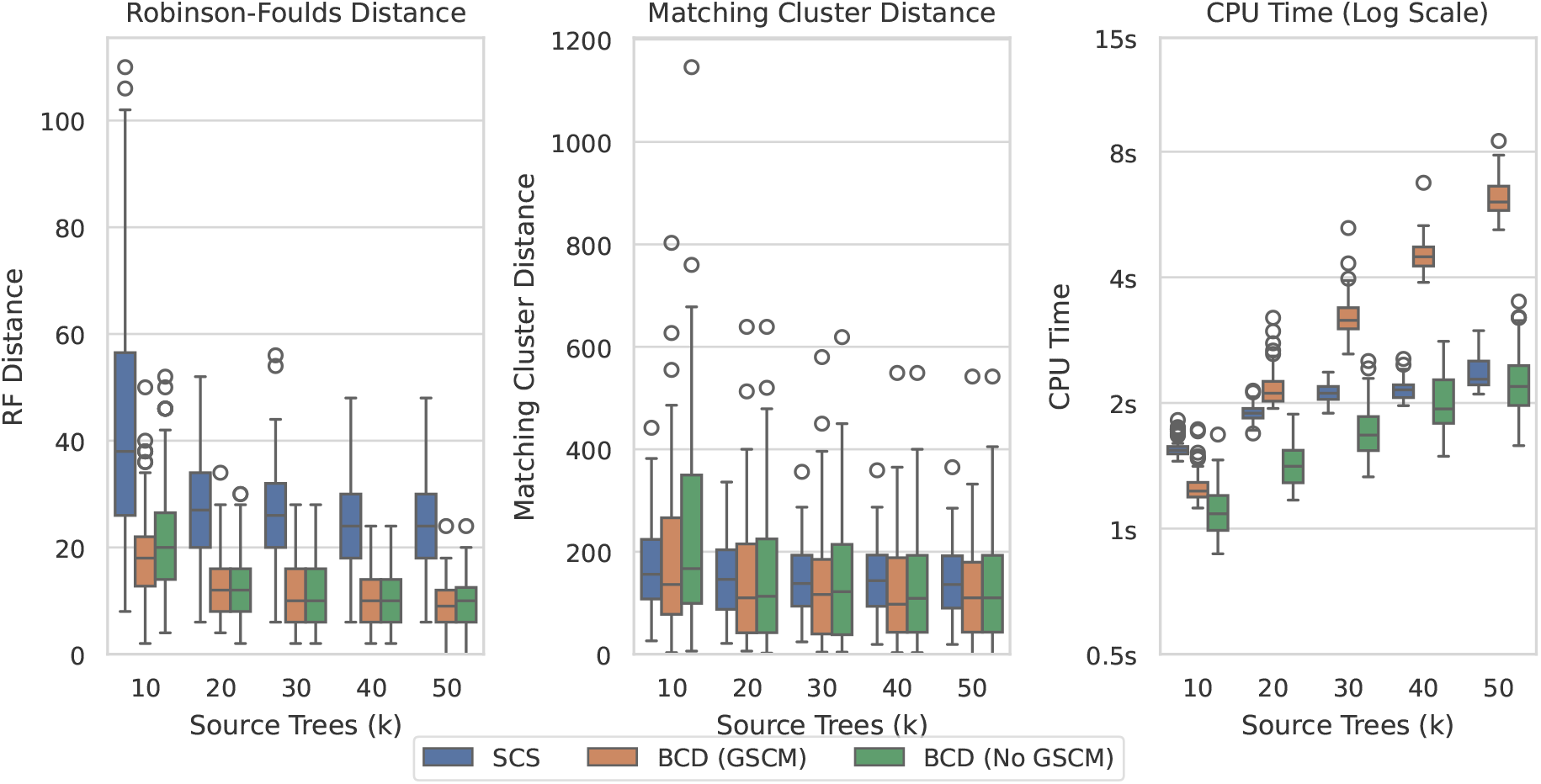
Spectral Cluster Supertree vs Bad Clade Deletion on the SuperTriplets dataset with a deletion rate of 25%.

**Figure A.8:**
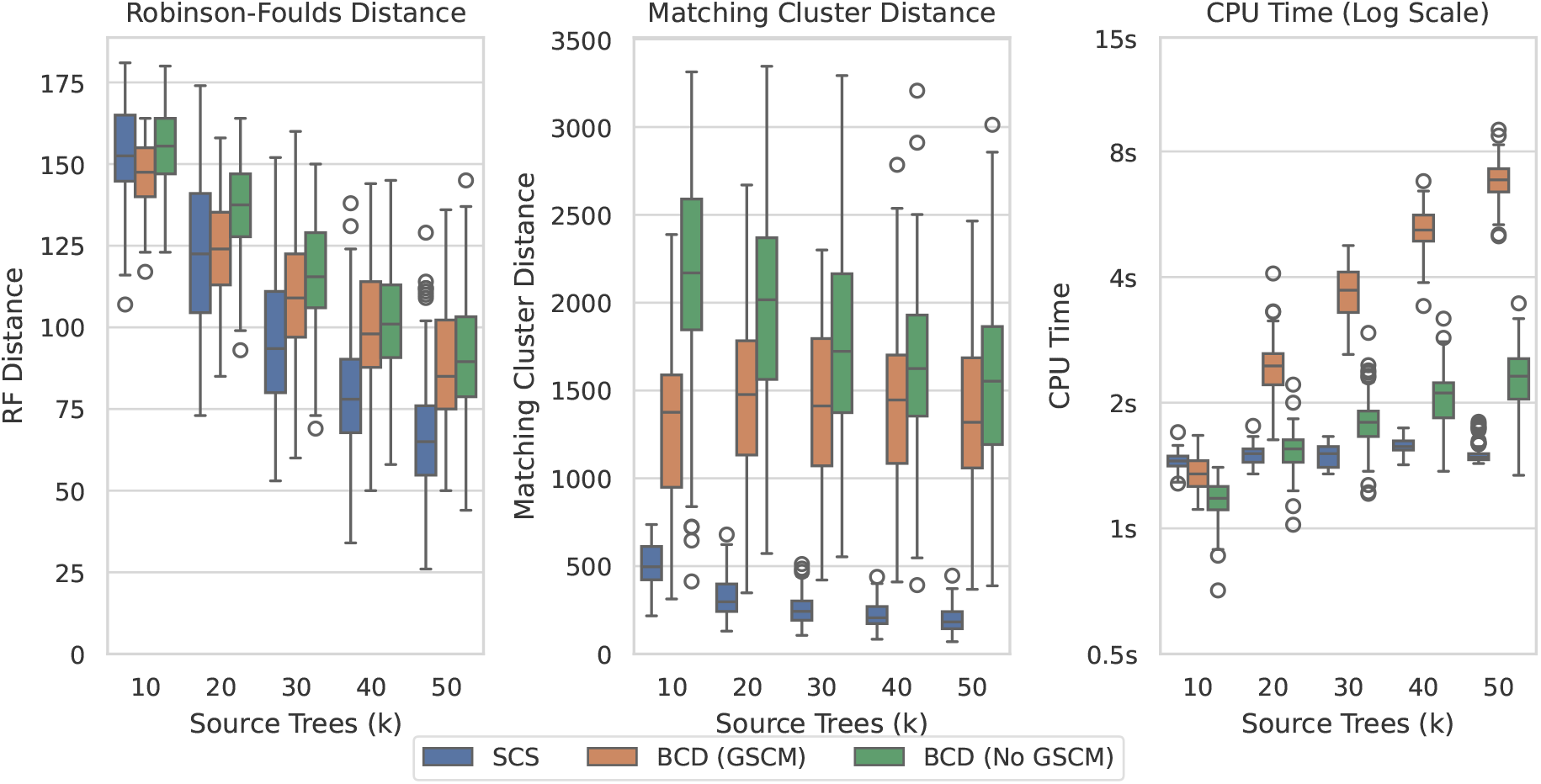
Spectral Cluster Supertree vs Bad Clade Deletion on the SuperTriplets dataset with a deletion rate of 75%.

