## Supplementary Materials for "Spectral Cluster Supertree: fast and statistically robust merging of rooted phylogenetic trees"

### SUPPLEMENTARY MATERIAL FOR: *Spectral Cluster Supertree: fast and statistically robust merging of rooted phylogenetic trees*

**Robert N. McArthur, Ahad N. Zehmakan, Michael A. Charleston and Gavin Huttley**

#### Overview

The supplementary material contains figures displaying the full results of Spectral Cluster Supertree (SCS) against Bad Clade Deletion (BCD) [3] over all datasets. Unlike the paper, it includes the $F_1$ score as defined there in each of the figures. It is notable however that there is a direct mapping between the $F_1$ score and the Robinson-Foulds [5] distance, as illustrated by the main paper. For the figures in this supplementary material, higher values are better only for the $F_1$ score. Lower values are better for all other graphs. Time results are shown on a logarithmic scale. The code used to generate these figures, as well as the raw results, has been archived online (https://doi.org/10.5281/zenodo.11118313).

#### SCS-DCM-IQ Dataset

Our SCS-DCM-IQ dataset was created to mimic what may be encountered by divide and conquer algorithms for phylogenetic reconstruction. Figures S1-S5 compare SCS to BCD with increasing amounts of taxa. While BCD (GSCM) tends to achieve more exactly correct clades (RF distance [5] and $F_1$ score [3]), the overall topological accuracy of SCS appears to be consistently superior (Matching Cluster distance [1]). SCS is also much faster than BCD, on the largest dataset taking on average ~20 seconds per problem, where BCD takes ~2 hours.

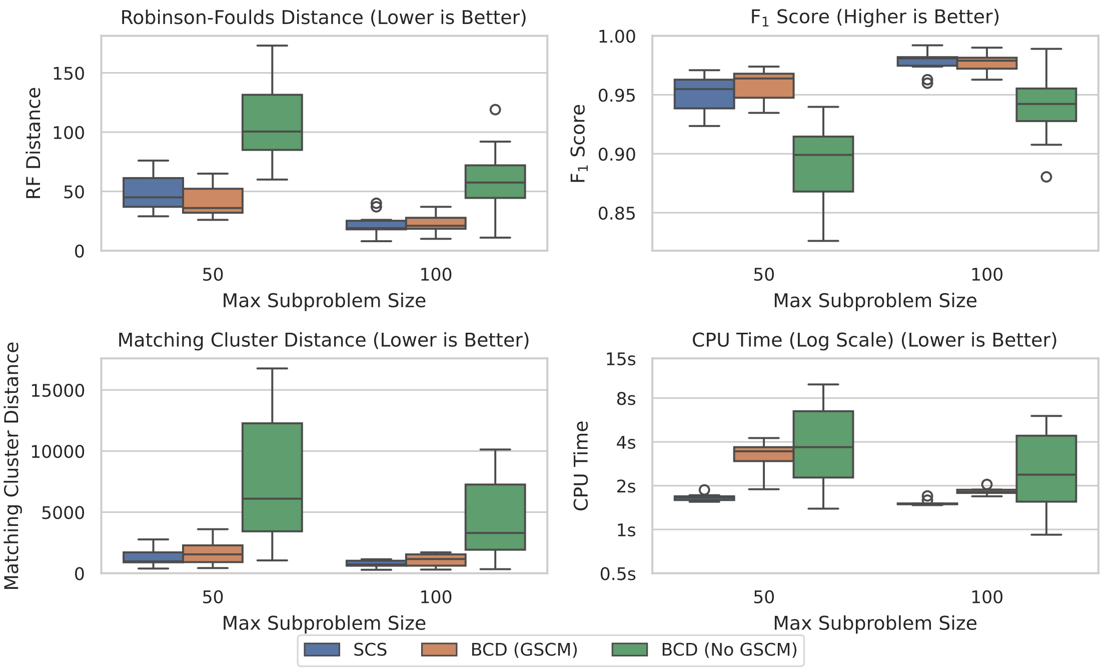


Figure S1: SCS vs BCD on the SCS-DCM-IQ dataset with 500 taxa. All methods solved all problems within the timeout.

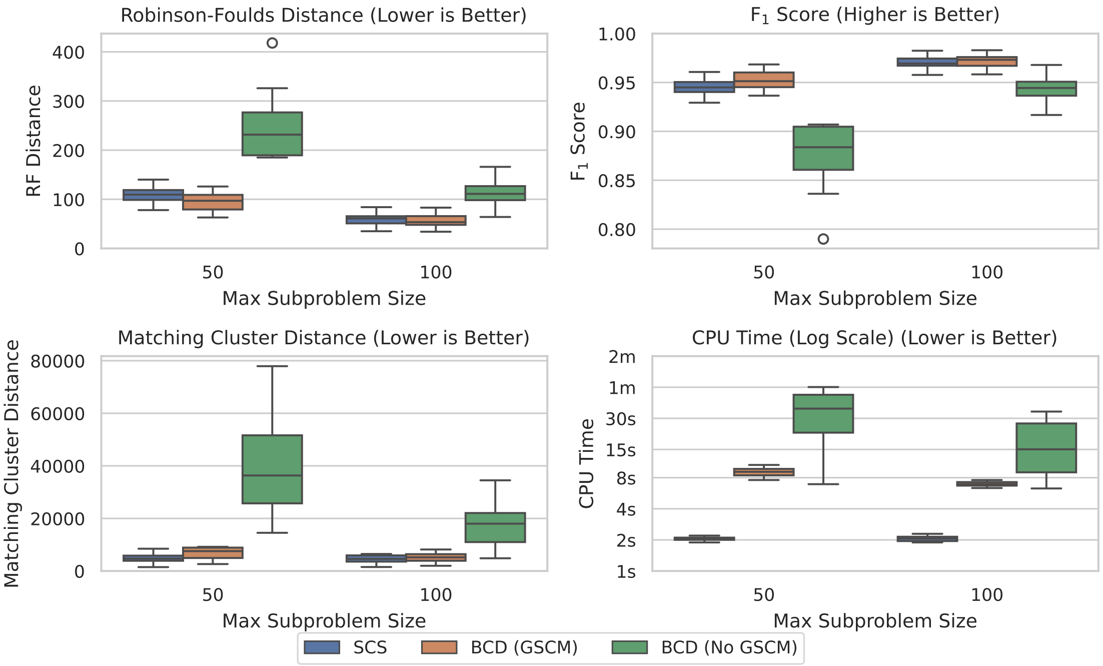


Figure S2: SCS vs BCD on the SCS-DCM-IQ dataset with 1000 taxa. All methods solved all problems within the timeout.

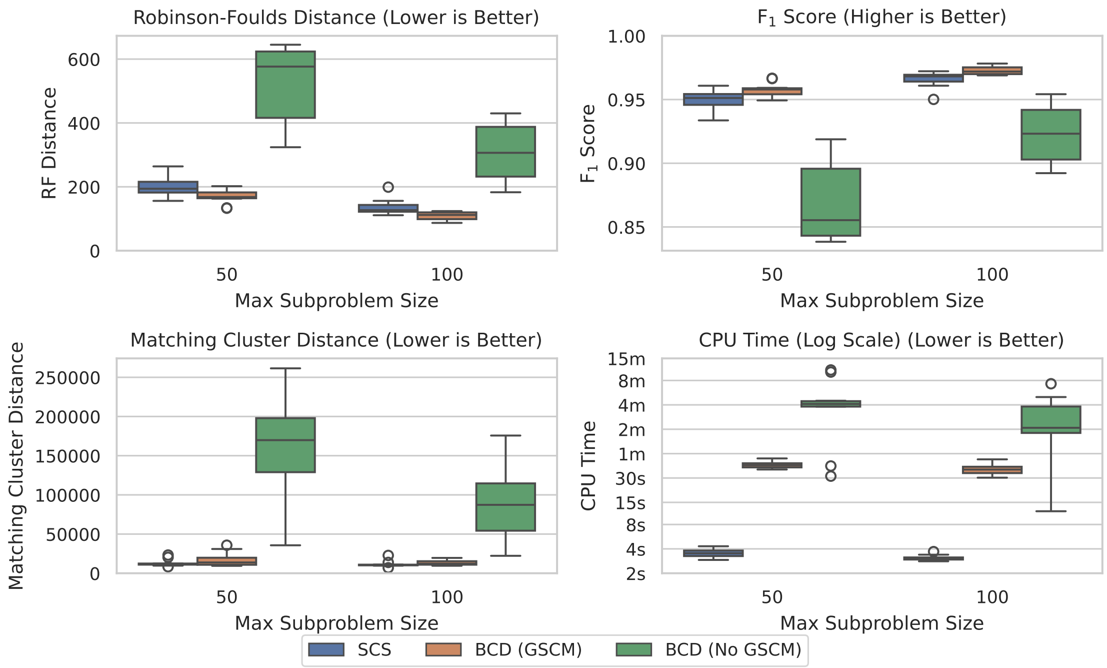


Figure S3: SCS vs BCD on the SCS-DCM-IQ dataset with 2000 taxa. All methods solved all problems within the timeout.

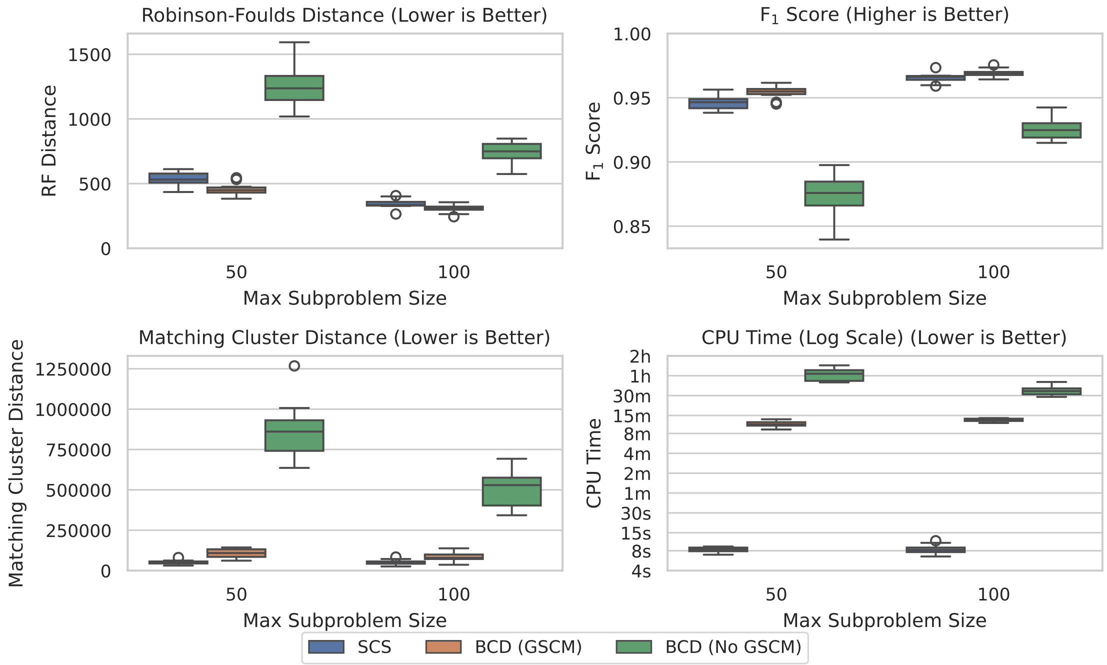


Figure S4: SCS vs BCD on the SCS-DCM-IQ dataset with 5000 taxa. All methods solved all problems within the timeout.

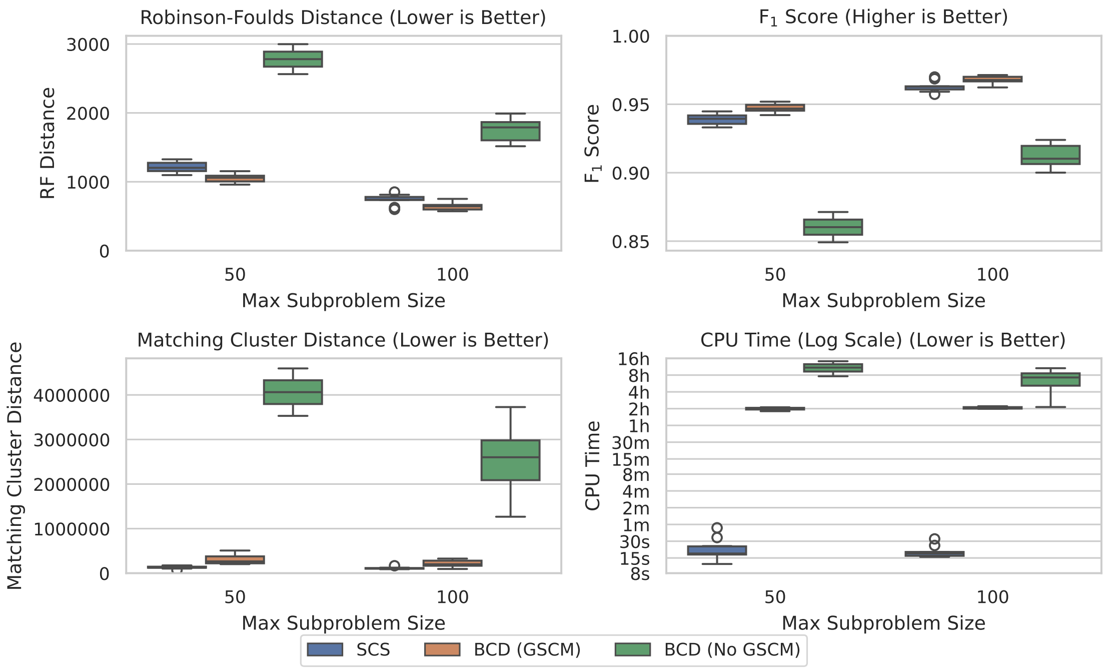


Figure S5: SCS vs BCD on the SCS-DCM-IQ dataset with 10000 taxa. BCD without GSCM processing only solved the first 2 and 6 problems within the timeout for the 50 and 100 max subproblem sizes respectively. The other methods solved all ten problems.

#### SMIDGenOG-5500 Dataset

The SMIDGenOG-5500 dataset [3] is a large scale dataset containing on average 5500 taxa. It follows the SMIDGen protocol [6], differing in how it generates scaffold trees due to the scale of the dataset. It aims to emulate what may be encountered by systematists at a large scale. Figure S6 compares SCS to BCD over this dataset. While BCD (GSCM) again achieves more exactly correct clades, the overall topological accuracy of the tree for SCS is vastly superior. SCS takes on the order of a couple of minutes to solve these problems, whereas BCD can take multiple hours.

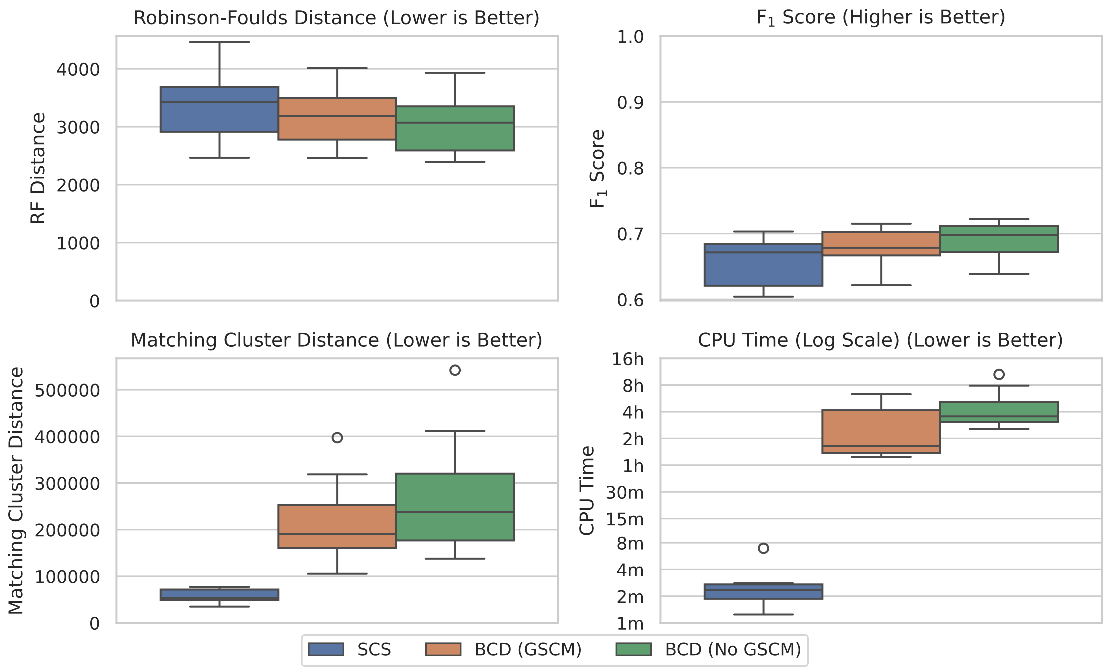


Figure S6: SCS vs BCD on the SMIDGenOG-5500 dataset. As there is no parameterisation for this dataset, no x-axis is displayed. All methods solved all problem instances within the timeout.

#### SMIDGenOG Dataset

The SMIDGenOG dataset [2] was developed using the SMIDGen protocol [6] in a rooted context. The datasets generated through the SMIDGen protocol aims to imitate data collection processes typically used by systematists. The dataset contains a scaffold tree sampling a percentage of the taxa, and many densely sampled clade based source trees. Figures S7-S9 compares SCS to BCD in increasing order of number of taxa. BCD (GSCM) appears to perform better than SCS over this dataset on all accuracy metrics, though the degree of improvement appears to decrease as the scaffold factor decreases. The time results are all small enough to not matter for practical purposes between SCS and BCD (GSCM).

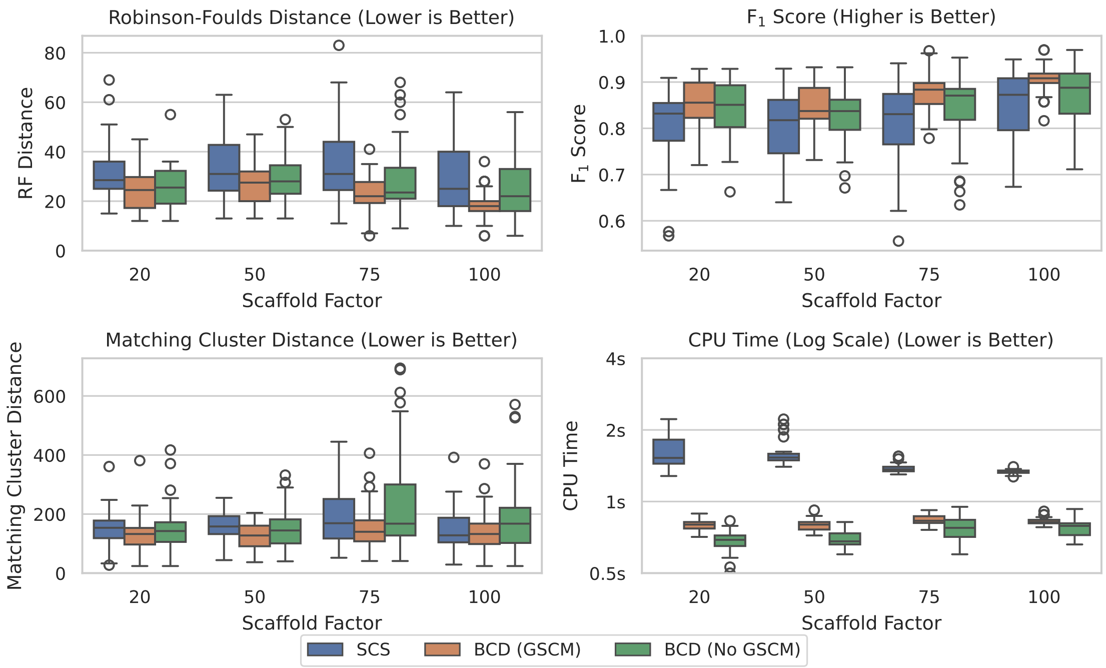


Figure S7: SCS vs BCD on the SMIDGenOG dataset with 100 taxa. The SMIDGenOG dataset contains one scaffold tree sampling over "Scaffold Factor" percent of the taxa, as well as many densely sampled clade-based source trees.

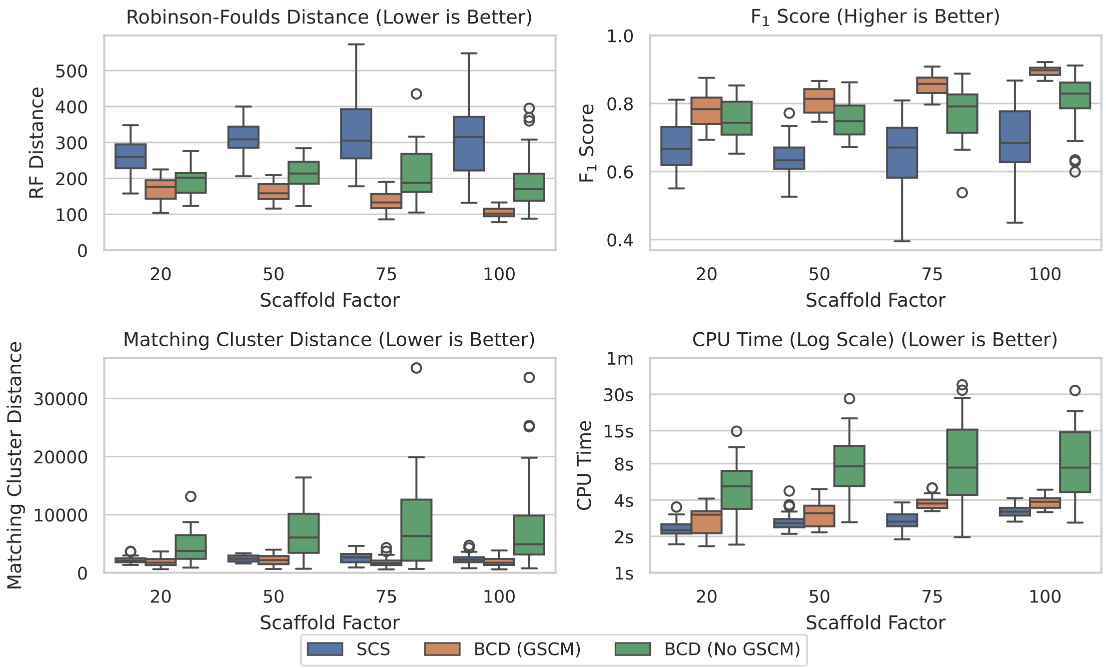


Figure S8: SCS vs BCD on the SMIDGenOG dataset with 500 taxa. The SMIDGenOG dataset contains one scaffold tree sampling over "Scaffold Factor" percent of the taxa, as well as many densely sampled clade-based source trees.

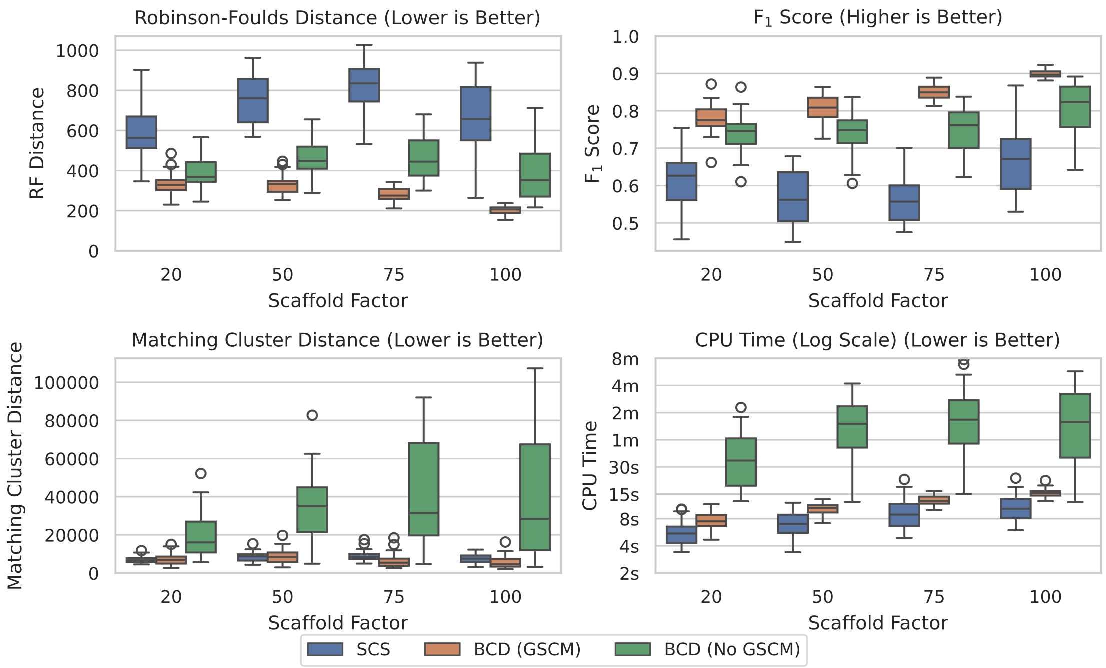


Figure S9: SCS vs BCD on the SMIDGenOG dataset with 1000 taxa. The SMIDGenOG dataset contains one scaffold tree sampling over “Scaffold Factor” percent of the taxa, as well as many densely sampled clade-based source trees.

#### SuperTriplets Dataset

The SuperTriplets dataset [4] explores the effect of the number of taxa present in each of the source trees after a percentage of them are removed, and the number of source trees, on supertree reconstruction accuracy. Figures S10-S12 compare BCD to SCS in increasing order of deletion rate. For the 25% deletion rate, SCS performs worse in terms of accuracy for the $\mathrm{RF}$ and $F_1$ metrics, though has a lower spread of values (albeit higher median) for the Matching Cluster distance. As the deletion rate of taxa increases (i.e. the number of taxa represented in each source tree decreases), the degree of improvement of SCS over BCD increases with respect to the Matching Cluster distance in particular. SCS even generally outperforms BCD in terms of the $\mathrm{RF}$ distance for the 75% deletion rate. The time results here are all low enough to not matter for practical purposes.

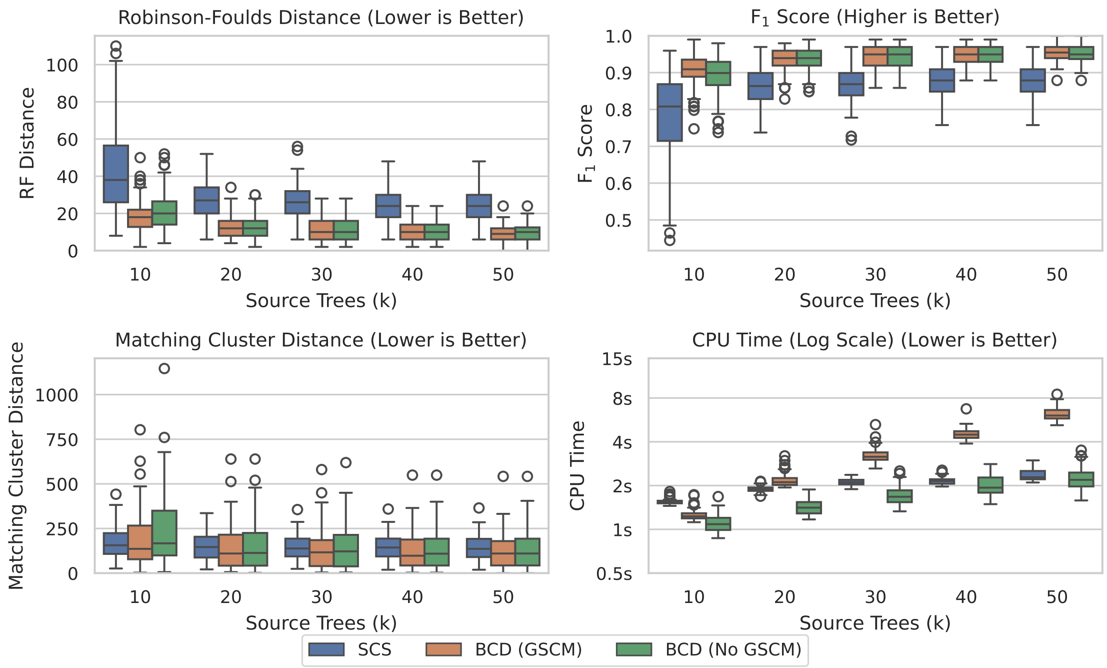


Figure S10: SCS vs BCD on the SuperTriplets dataset with a deletion rate of 25%.

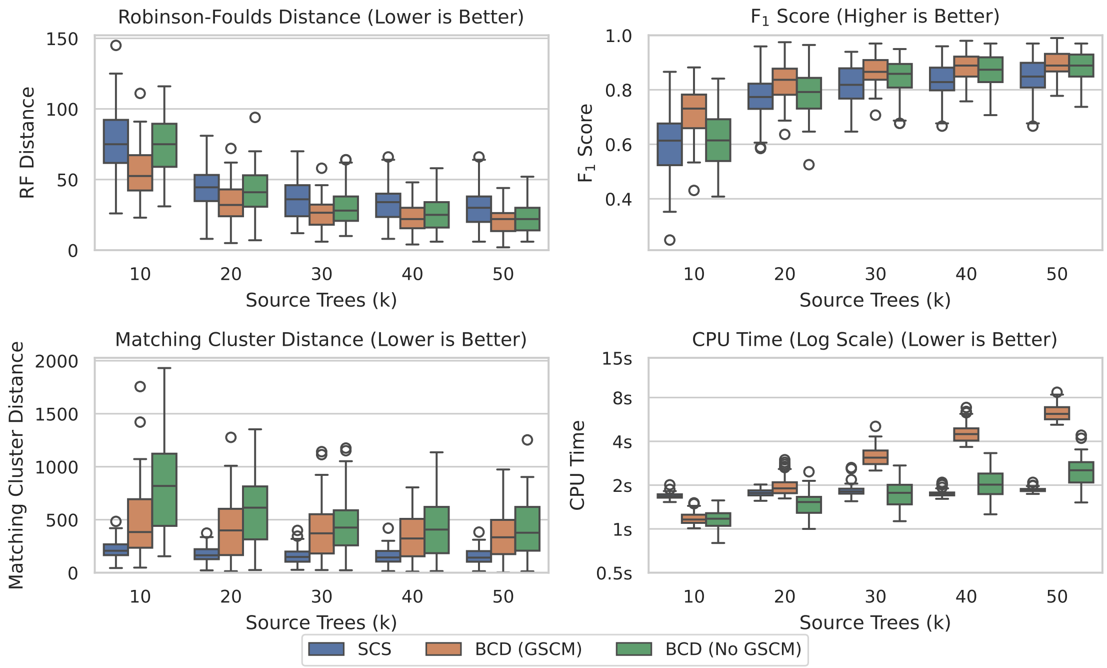


Figure S11: SCS vs BCD on the SuperTriplets dataset with a deletion rate of 50%.

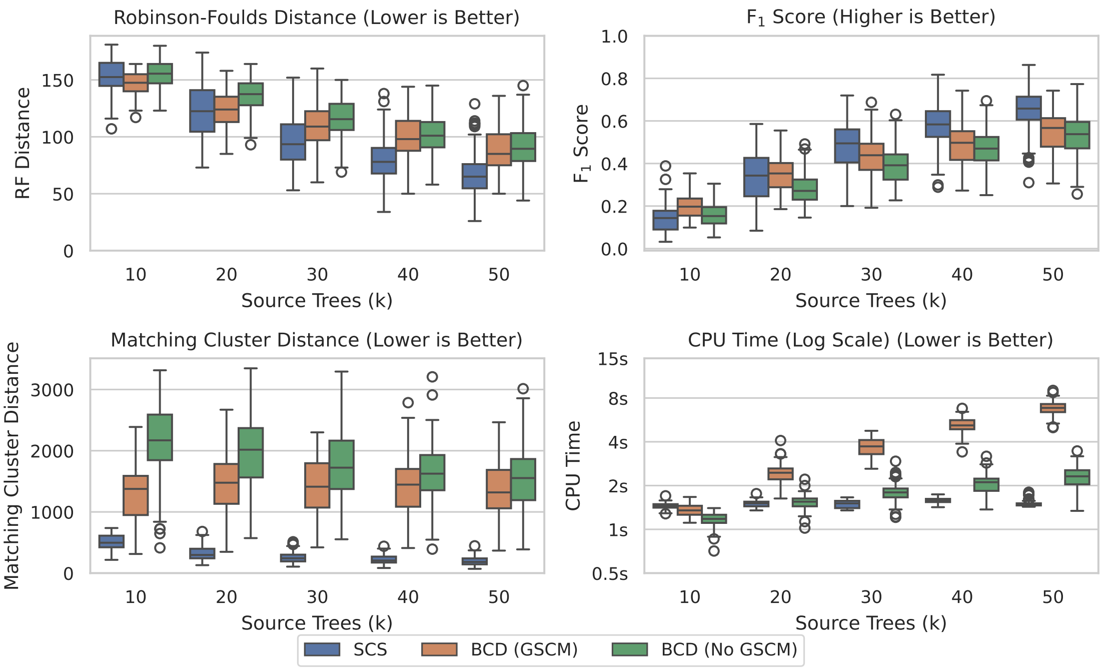


Figure S12: SCS vs BCD on the SuperTriplets dataset with a deletion rate of 75%.
